# Borg extrachromosomal elements of methane-oxidizing archaea have conserved and expressed genetic repertoires

**DOI:** 10.1101/2023.08.01.549754

**Authors:** Marie C. Schoelmerich, Lynn Ly, Jacob West-Roberts, Ling-Dong Shi, Cong Shen, Nikhil S. Malvankar, Najwa Taib, Simonetta Gribaldo, Ben J. Woodcroft, Christopher W. Schadt, Basem Al-Shayeb, Xiaoguang Dai, Christopher Mozsary, Scott Hickey, Christine He, John Beaulaurier, Sissel Juul, Rohan Sachdeva, Jillian F. Banfield

## Abstract

Borgs are huge extrachromosomal elements (ECE) of anaerobic methane-consuming “*Candidatus* Methanoperedens” archaea. Here, we used nanopore sequencing to validate published complete genomes curated from short reads and to reconstruct new genomes. 13 complete and four near-complete linear genomes share 40 genes that define a largely syntenous genome backbone. We use these conserved genes to identify new Borgs from peatland soil and to delineate Borg phylogeny, revealing two major clades. Remarkably, Borg genes encoding OmcZ nanowire-like electron-exporting cytochromes and cell surface proteins are more highly expressed than those of host *Methanoperedens*, indicating that Borgs augment the *Methanoperedens* activity *in situ*. We reconstructed the first complete 4.00 Mbp genome for a *Methanoperedens* that is inferred to be a Borg host and predicted its methylation motifs, which differ from pervasive TC and CC methylation motifs of the Borgs. Thus, methylation may enable *Methanoperedens* to distinguish their genomes from those of Borgs. Very high Borg to *Methanoperedens* ratios and structural predictions suggest that Borgs may be capable of encapsulation. The findings clearly define Borgs as a distinct class of ECE with shared genomic signatures, establish their diversification from a common ancestor with genetic inheritance, and raise the possibility of periodic existence outside of host cells.

## Introduction

Of all genetic entities in the biosphere, extrachromosomal elements of archaea may be the least well understood. Existing databases of archaeal viruses, plasmids and mini-chromosomes are limited (NCBI Virus, 556 viruses, Jun-02-2023 and NCBI Plasmids, 481 plasmids; www.ncbi.nlm.nih.gov/genome/browse#!/overview/) and generally taxonomically restricted to better known archaeal groups, such as the orders Sulfolobales and Halobacteriales. For many major groups of archaea not even a single ECE has been described and those that affiliate with uncultivated archaea are only inferred from metagenomic sequence information.

We recently described a seemingly new type of archaeal ECE that is not classifiable as virus or plasmid ^1^. They are unusually large, up to 1.1 Mbp in length, and have linear genomes terminated by kilobase-scale long inverted terminal repeats (ITR). The genes are encoded on two replichores of very uneven length, with essentially all genes on each replichore carried on a single strand. These ECEs are predicted to replicate in *Methanoperedens* archaea based on sequence similarity, phylogeny, shared abundance-based co-occurrence patterns and CRISPR targeting ^1^. Because of their propensity to assimilate large numbers of genes from organisms, especially but not limited to their host *Methanoperedens*, they were named “Borgs” ^1^. To date, their sequences have only been recovered from environmental metagenomic datasets from terrestrial subsurface ecosystems and interestingly often encode components of key metabolic processes such as anaerobic methane oxidation and nitrogen fixation ^1^.

When new and unusual ECEs are recovered from metagenomic datasets, they may be questioned because methods for curation of short read assemblies into complete genomes are poorly understood and rarely undertaken. Curation is important because short read assemblies are often fragmented and without a complete genome it is impossible to rule out the possibility that the sequence is, for example, a virus for which the structural proteins were not recovered, or an ECE that is integrated (e.g., pro-plasmid). Lack of complete or near-complete genomes also precludes identification of genes that are universally present and comparison of genome architectures. Here, we used long read sequences from Oxford Nanopore Technologies that can span repeated regions to validate the overall topologies of some of our short read-derived complete Borg genomes. Nanopore-derived sequence information was combined with information from more accurate and deeply sequenced short Illumina reads and contigs to reconstruct additional complete and near-complete genomes.

Anaerobic methane-oxidizing archaea of the genus *Methanoperedens* are known primarily from bioreactor co-cultures ^2, 3^. To date, only one complete *Methanoperedens* genome has been reported ^4^ and it does not host Borgs. We obtained one complete and one near complete genome for two *Methanoperedens* which are both predicted to host Borgs. These genomes enabled comparison of Borg and host genetic repertoires as well as gene expression patterns. Overall, our results greatly increase sampling of the genomes and genetic repertoires of Borgs, establish that previously reported Borg characteristics are conserved, and reveal a mostly syntenous set of conserved marker genes that could be used to identify new Borgs and define their phylogeny.

## Results

### Evaluation of existing complete Borg genome sequences

The majority of previously reported Borg genomes derived from soil sampled from a seasonal wetland in Lake County, northern California, and all were reconstructed using Illumina sequences ^1, 5^. We performed long read nanopore sequencing on a subset of the same samples to evaluate the accuracy of the manually curated Illumina-based complete genomes reconstructed from these samples and to recover new Borg genomes. We sequenced both native DNA and DNA amplified with multiple displacement amplification (MDA), which served as a negative control for methylation calling and had more accurate basecalls, due to fewer signal current disruptions caused by modified bases. Obviously erroneously assembled regions, such as chimeric joins that were often accompanied by a substantial jump in GC content and mirror image artifacts that resulted from the multiple displacement amplification process were removed. The corrected nanopore-assembled sequences validate the genomes of Green, Ochre, Purple, Orange, and Black Borgs **(Extended Data Figure 1 a-d, Extended Data Table 1**). In most cases the near complete curated nanopore-derived sequences extend into, thus confirming, the ITR.

The ∼1.1 Mbp Green Borg is the largest Borg genome recovered to date. After removal of an obviously chimeric start (not supported by mapped Illumina reads), the nanopore genome aligns completely with the published Illumina-based genome (**Extended Data Figure 1a**). The Green Borg genome was reported to have a slightly more complex genome architecture than all other Borgs due to a switch in coding strand over a small region of the large replichore (**Supplementary Figure 1**). This is verified by the nanopore assembly (**Extended Data Figure 1a**). Essentially all disagreements with the published genome involved a larger than expected number of repeat units in the tandem repeat (TR) regions. This is unsurprising, as some TR regions are longer than the Illumina read lengths. However, the number of TR units in TR regions is often variable within populations ^5^, so a combination of deep Illumina and some longer nanopore reads may be best used to define the TR regions.

### Improved and new Borg genomes reveal conserved features

We used nanopore read assemblies to improve the quality of known Borg genomes and to reconstruct new genomes. Nanopore reads were assembled, mirror image sequence blocks and chimeras were removed and then sequencing errors were corrected by automated short- and long-read polishing methods followed by careful manual curation using Illumina reads and assembled Illumina contigs. Removal of single base pair errors, such as in homopolymers, is important as these can interrupt gene predictions. Ultimately, we reconstructed two new complete Borg genomes (Amethyst and Amber) and one essentially complete genome (Cobalt Borg, curated to 1.08 Mbp and into the ITR) (**Extended Data Figure 1e**). In addition, four new near-complete genomes for Iris, Emerald, Ruby and Viridian Borgs were curated into single long contigs. Emerald and Viridian genomes include both replichores (**Extended Data Table 1, Supplementary Figure 2**).

Amethyst Borg is related to Blue Borg, the genome of which is currently available only in draft form. Borg-like contigs with consistently high coverage and low GC content were joined and aligned to the new Amethyst genome to generate a concatenated ∼876 kbp Blue Borg sequence comprising the expected single strand coding of two replichores and termination by ITR.

All Borg genomes are linear and have long ITR, with an overall low GC content (∼32-35%). However, there are local regions with elevated GC content. Some of these have high sequence similarity and very similar GC content to genes of *Methanoperedens*. There is near perfect single strand coding on each of the replichores, and many of the encoded proteins are specific to Borgs. The majority of genes (63%) lack a taxonomic affiliation (12,875 of 20,469). On average, 19.6% were taxonomically classified as archaeal (**Supplementary Table 1**). All newly reported genomes have the expected GC skew, consistent with replication initiation from the termini (**Supplementary Figure 1**).

All Borg genomes harbor pervasive nucleotide TR that are located in intergenic regions and within open reading frames (**Extended Data Table 1**). Approximately half of these TRs occur in genes, where they essentially always introduce amino acid repeats that generally confer local intrinsic disorder, as reported by us previously ^5^. Representative examples in Amber and Amethyst Borg are TRs located in multiheme cytochromes (MHC) (**Supplementary Figure 3**), which often harbor aaTRs ^5^. These regions may provide sites for post translational modifications, mediate complex formation or bind small molecules ^5, 6^. Outstanding is Emerald Borg with only one TR in an intergenic region, and eighteen within ORFs. Overall, the well defined Borg genomes have consistent features, supporting their assignment to a specific class of extrachromosomal elements.

### Borgs have a core genome that suggests a shared evolutionary origin

We used the 17 complete and near-complete genomes (**Extended Data Table 1**) to identify conserved genes. Protein family clustering of all 20,280 Borg proteins revealed that 88.0% of Borg proteins have at least one homolog in another Borg (**Supplementary Table 2**). A comparison of shared protein subfamilies revealed that Black and Brown Borg share the highest number of protein subfamilies, and Emerald and Red Borgs are most distantly related (**Extended Data Figure 2a,b**).

Of the 2,598 Borg protein subfamilies, 107 are present in all Borgs and thus make up the core Borg proteome (**Supplementary Table 2, Supplementary Table 3**). This core encompasses 40 subfamilies representing marker proteins (n=1 in each Borg), 22 subfamilies representing near marker proteins (n=1 in 16/17 Borgs), and 45 subfamilies representing multicopy marker proteins (multiple homologs in each Borg). Only ∼18% (6/40) of the Borg marker proteins have homologs in the complete *Methanoperedens* genome.

Mapping the 40 Borg marker genes onto each Borg genome revealed a fairly consistent localization of each subfamily gene across the entire large replichore, implying a shared genomic backbone (**Figure 1**). This analysis also demonstrated that subsets of conserved genes are co-localized, suggesting a functional relationship and motivating a more general analysis of shared Borg proteins.

**Figure 1.**
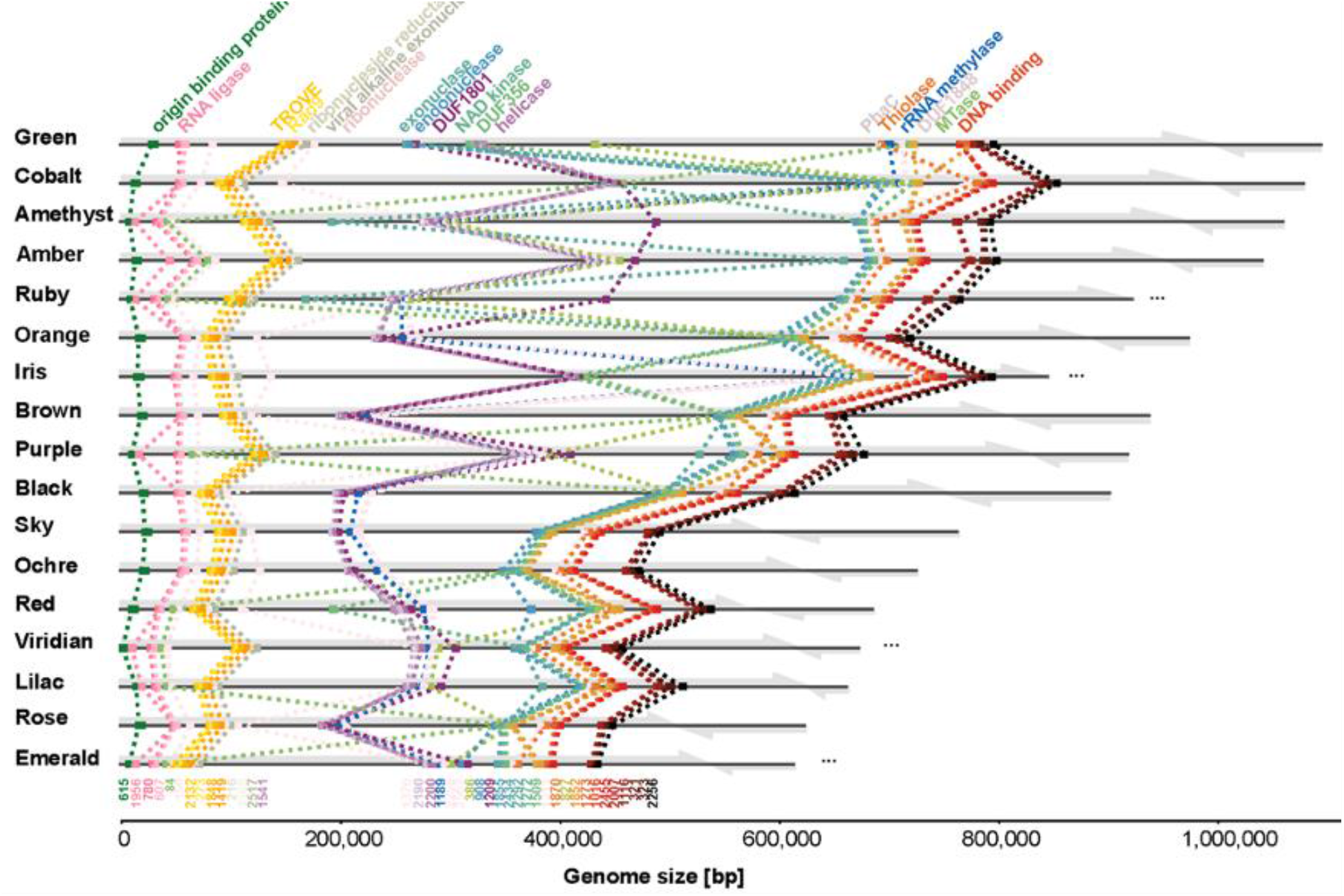
Borg protein family clustering defines relatedness and reveals syntenous localization of marker genes on Borg genomes. Complete Borg genomes are listed in order of decreasing length, with near-complete genomes included in order of the assembled lengths of the large replichore. Each replichore is represented by a gray arrow. Dots indicate the locations of the 40 marker genes (n=1 in every Borg). Functional annotations are shown on the top and subfamily numbers on the bottom of the figure.

### Core Borg proteins for replication, nucleotide processing, cell decoration, signaling, carbon metabolism and redox

To investigate the hypothesis that co-localized genes are functionally related we grouped together marker, near marker, and multicopy proteins based on colocalization, considering these core proteins to be co-localized if they occur within ten genes in a minimum of five Borg genomes. This generated eight core clusters comprising 84 of the 107 core subfamilies (**Extended Data Figure 3a**). We then examined the putative functions of the encoded core proteins, leveraging a combination of sequence-based functional annotations and structure-based (AlphaFold2 (AF2) ^7^) homology predictions (**Extended Data Figure 3b**). We found structural matches for five core proteins that did not have any sequence-based annotations. One of these matches the DarG toxin (subfam2258, PDB match 5M3E) of the toxin-antitoxin system DarTG. DarG expression affects DNA replication by catalyzing the reversible ADP-ribosylation of thymidines in ssDNA ^8^. We speculate that this protein could be involved in regulating Borg or host replication. The other four have credible matches to viral capsid proteins (see below).

Some core proteins are predicted to be involved in replicating Borg genomes. The 18 (including Blue) Borg genomes have a marker protein encoded relatively close to the start of the large replichore that has functional domains that match those of the Herpesvirus OBP (PF02399, Herpes_ori_bp). OBP binds to the origin of replication and initiates the formation of a pre-replication complex ^9^. Emerald is the first reported Borg to encode a typical replication-initiation protein, cdc6, situated just five genes downstream from the OBP. The second core gene cluster encoded on Borg genomes (green in **Extended Data Figure 3**) includes a DNA polymerase B, accompanied by a YspA-related gene (subfam0180 and subfam1707) that could function as a sensor of nucleotide, nucleotide-derived ligands, or nucleic acids ^10^. The DNA polymerase is highly conserved in Borgs and likely involved in replication.

The co-localized core proteome encodes many genes whose predicted functions imply roles in sensing and responding to DNA damage. For example, the second core gene cluster includes an RNA ligase capable of repairing breaks in nicked DNA:RNA and RNA:RNA. Interestingly, the predicted Borg protein structure is most akin to the RNA ligase from *Escherichia* virus T4 (PDB match 1s68, RMSD = 2.72), known for repairing tRNA breaks induced by a host protein following phage infection ^11^. This may indicate the need for Borgs to protect their tRNA (5-23 tRNAs per Borg genome) and the host tRNA from degradation by the host archaeon. This genomic context also includes a radical SAM protein, a homing endonuclease, and a ClpP protease (PDB match 5vz2) which could be degrading defective and misfolded proteins; altered function of this protein has been shown to affect virulence and infectivity of pathogens ^12^.

The third (orange in **Extended Data Figure 3**) cluster features a TROVE domain protein, likely involved in RNA-binding and degradation ^13^, and Rad9, which may monitor and respond to DNA damage. Following this is a cluster potentially involved in DNA processing/modification (**Supplementary Figure 4**), including a DNA methylase (subfam0757 and subfam0999 where the DNA methylase is fused to an intein) that may introduce N^4^ cytosine-specific or N^6^ adenine-specific DNA methylations (4mC or 6mA). No close homologs of this methylase exist in any *Methanoperedens* (highest amino acid identity is 51%).

Further suggesting roles in sensing and responding to damage, the cluster represented in blue in **Extended Data Figure 3** includes a protein (21) with remarkable structural homology to the 3’-5’ DNA exonuclease Cap18 from *Escherichia coli* (PDB match 7t2s, RMSD = 0.96) ^14^. This is part of the bacterial immune cyclic oligonucleotide-based anti-phage signaling (CBASS) system, yet no other part of the system has been identified in Borgs and *Methanoperedens*. Also present is an endonuclease III (24, PDB match 1p59, RMSD = 1.66) that may function as a DNA glycosylase with repair activity specific for oxidized pyrimidine lesions in duplex DNA ^15^. Other small clusters encode a putative Pnkp1–Rnl–Hen1 RNA repair complex (pink clusters) which is required to repair ribotoxin-cleaved RNA ^16^, a manganese catalase potentially involved in hydrogen peroxide detoxification (brown clusters), or cell decoration machinery (purple cluster and members of the orange cluster). One cluster (red in **Extended Data Figure 3**) comprises seven protein subfamilies which encode a Rad3-related helicase (15), capable of unwinding dsDNA or DNA:RNA ^17^, and possibly contributing to nucleotide excision repair, and an orotate phosphoribosyltransferase (OPRTase), putatively involved in pyrimidine synthesis.

A subset of the core families are predicted to have roles in metabolism. Strikingly, all Borgs encode the polyhydroxyalkanoate (PHA) pathway (blue in **Extended Data Figure 3**) that likely functions in PHA degradation, given the domain annotations and structural homology to a hydrolase (PhaC, PDB match 3om8). The PHA pathway also exists in many *Methanoperedens* genomes and has been implicated in PHA synthesis ^4^.

Nine Borgs encode a *nif* gene cluster with elevated GC content and sequence similarity to clusters in *Methanoperedens*. In Green, Ochre and Sky Borgs and *Methanoperedens* the *nif* cluster is located far distant from the PHA cluster. In Cobalt, Iris and Orange Borgs it interrupts the PHA cluster, and in Purple, Amethyst, and Amber Borg genomes the *nif* cluster is located just downstream of the PHA cluster (**Extended Data Figure 4**). The colocalization of nitrogenase and PHA clusters suggests that they could be functionally connected. We speculate that the reducing power (NAD(P)H) from PHA degradation could be harnessed by nitrogenase to fuel nitrogen fixation and potentially even concomitant H_2_ formation ^18^. Also within this cluster is an NAD kinase, which regulates cellular NADP(H) levels, a crucial redox carrier in the PHA pathway. The NifH protein phylogeny groups the Amethyst sequence within one clade of *Methanoperedens* sequences and the Ochre sequence within another clade of *Methanoperedens* sequences, whereas most other Borg NifH form a Borg-specific clade. Elsewhere, Green, and Black Borgs encode an isolated NifH-like protein closely related to sequences found in *Methanoperedens*. These patterns point to repeated lateral transfer of *nif* genes into Borgs and sequences grouping with those of *Methanoperedens* may suggest Borg-host associations.

We identified 207 Borg-encoded multiheme cytochromes (MHCs that have ≥ 3 CxxCH) with up to 33 classical heme-binding motifs (**Supplementary Table 4**). Interestingly, MHCs represent one of the largest multicopy subfamilies (subfam1158, subfam1369, subfam2060, subfam2491), but given their very divergent sequences, only one subfamily emerged in the protein network analysis (gray in **Extended Data Figure 3**). Cobalt Borg encodes the maximum of 19 different MHCs. The Borg-encoded MHCs likely augment the ability of *Methanoperedens* to dispose of electrons produced during methane oxidation.

### Soil distribution patterns as a clue to *Methanoperedens* - Borg linkages

The pairing of Borgs and hosts is a topic we attempted to address experimentally without success (see **Methods**). However, Borg and *Methanoperedens* relative abundances increase with soil depth and Borg and *Methanoperedens* types vary with soil depth (e.g., Rose in shallower soils; Black and Brown in medium depth soils, Viridian in the deeper soils; **Extended Data Figure 5**), providing a clue to Borg - host associations. For example, when organized by presence but without strong reliance on abundance data, the patterns support the previous prediction that Black Borg replicates in *Methanperedens* named bMp, and hints that Orange and Ochre Borgs replicate in another *Methanoperedens* type, cMp. Supporting the Orange host prediction, the Orange Borg exhibits more homologs (21%, 258/1,243) with cMp than other Borgs (**Supplementary Table 5**).

We used the abundance data to seek statistically significant correlations and found support for the replication of Black Borg in bMp. Interestingly, Brown Borg abundances are correlated with those of Black Borg, suggesting that it may also replicate in bMp. These Borgs also show significant correlations (R^2^>0.995) with five additional *Methanoperedens* species. A weaker correlation (R^2^=0.961) was identified for Sky Borg and *Methanoperedens*_80cm_43_82 (**Supplementary Table 6**). The lack of strong correlations for the majority of Borgs from the wetland may be due to (a) variable Borg to *Methanoperedens* genome copy number (as suggested previously; ^1^), (b) the same Borg resident in multiple different hosts, or (c) Borg DNA derived from outside of *Methanoperedens* cells.

Clarifying these possibilities, we note that Orange Borgs is 72 times more abundant than the most abundant co-occurring *Methanoperedens* in 1.05 m deep soil (**Extended Data Figure 5, Supplementary Table 7**). Such a high copy number seems very unlikely, especially if there is a stable association, for example this Borg existing as a plasmid-like element in a single host. Even if Orange Borg was present in every single type of coexisting *Methanoperedens* cell, the abundance ratio is still 7.5 to 1. This also seems unlikely, especially given two other abundant Borgs (Ochre and Cobalt) co-occur in this sample. Simply comparing the totals for all Borgs and all *Methanoperedens* in the 1.05 m sample yields a predicted average genome copy number ratio of 8.8 to 1. These observations raise the possibility that some Borg DNA derives from cell-free genomes. Given no strong indication of DNA degradation from the ends of their linear genomes (based on read mapping), we consider the possibility that the Borg DNA can exist external to the cell, and that it is protected, possibly via encapsulation.

### Borgs encode a putative capsid and replication machinery akin to eukaryotic viruses

The AF2-generated protein structure prediction analysis suggested that four core proteins without any sequence-inferred annotations had best structural matches with capsid proteins from *Haloarcula hispanica* virus SH1 (PDB 6qt9), and from the archaeal extremophilic internal membrane-containing *Haloarcula californiae* icosahedral virus 1 (HCIV-1, PDB 6h9c) (**Extended Data Figure 6, Supplementary Table 8**) ^19, 20^. Like Borgs, these viruses have linear dsDNA genomes terminated by inverted repeats, yet they are only ∼30 kbp. The Borg-encoded putative capsid proteins in turn open the possibility of Borg encapsulation at a point in their existence.

CheckV ^21^ (v1.0.1) detected between 11 to 21 viral-like genes per Borg genome (**Supplementary Table 9**) whereas geNomad ^22^ (v1.5.1) detected 39 possible provirus regions and no plasmid genes (**Supplementary Table 10**). One credible match found in all Borgs and many linear viruses is the Borg marker subfam2517. This protein functionally and structurally matches a viral recombinase/alkaline exonuclease (**Extended Data Table 2, Supplementary Figure 4**), which is crucial in recombination and ultimately replication of viruses with linear dsDNA ^23^. Manual inspection of the annotated Borg proteins revealed that 48 individual Borg proteins, including a marker protein (subfam0386), are annotated as related to KSHV latency-associated nuclear antigen, which promotes Herpesvirus persistence ^24^. Interestingly, a protein with similar domain annotations (Herpes_alk_exo) is involved in the replication of eukaryotic Herpesviruses (HSV) and required for the production of viral DNA ^25^. HSV encodes seven essential replication proteins, three of which may correspond to the Borg marker proteins DNA PolB and the PCNA-like Rad9, as well as a marker helicase C (**Extended Data Table 2**). All Borgs encode the OBP replication initiation protein similar to that from HSV. Moreover, all Borgs encode one copy of a tubulin-like protein (subfam2487). Similar cytoskeletal proteins in HSV form nanotubes that facilitate cell-to-cell contacts, and are essential for efficient viral spreading and replication ^26^. Notably, HSV has a double stranded DNA genome, some of which are linear, and many genomes (≥ 92) feature numerous tandem repeats. Together, these observations indicate surprising shared features with this (much smaller) eukaryotic virus type, and bring to light indications that Borgs share substantial replication related machinery with diverse viruses.

### Borgs and hosts have distinct DNA modifications

We used Illumina reads to correct a small number of base errors in a nanopore assembled sequence and generated a circular, complete 4,003,944 Mbp *Methanoperedens* genome (cMp). This genome is substantially larger than the only other complete ∼3.3 Mbp genome of *M. nitroreducens* ^4^, which does not host Borgs ^27^. The cMp genome is predicted to host Orange and Ochre Borgs (**Extended Data Figure 5**), and we also detected Black, Purple, Cobalt and Ruby Borgs in this sample (SR-VP_9_9_2021_72_4B_1.05m).

From the same soil sample we also reconstructed a near complete, 2.43 Mbp *Methanoperedens* genome (bMp) that is on four contigs. This genome was previously suggested to represent the host for Black Borg based on statistically significant abundance-based co-occurrence patterns (with Borg : host abundance ratios ranging from 0.5 to 8 x; ^1^).

Given abundance ratio variation and some very large predicted Borg copies per most abundant potential host genome, as well as tentative indications of viral proteins and genes involved in encapsulation, we hypothesized that Borgs and *Methanoperedens* have distinct methylation patterns to distinguish between self and non-self DNA. Thus, we searched for DNA methylation motifs in the nanopore datasets by comparing the mean current of native DNA vs. MDA amplified DNA using Nanodisco. This brought to light nine methylation motifs in the *Methanoperedens* genome, six of which were substantiated by read-level methylation calling with Megalodon using its models for 6mA and 5mC detection (**Figure 2a, Extended Data Table 3**). Performing the same analysis on the genomes of Orange, Ochre, Black, and Purple Borg revealed that they have a modification motif composed of YC, where Y stands for either pyrimidine T or C. Surprisingly, these motifs are found 184-192 times per 1 kbp of Borg genome, and they are particularly dominant (71%) on the non-coding strand of each replichore. Modification of cytosine residues was previously detected in the 72 kbp linear dsDNA genome of the STSV1 virus infecting *Sulfolobus tengchongensis* where it is proposed to protect the viral DNA from degradation through host machinery, and ensure selective regulation of viral replication and transcription ^28^. The YC modification motif is also reminiscent of CpG that is found in eukaryotes, including DNA viruses infecting eukaryotes, where it is important in regulating gene expression, but also serves an intricate regulatory role during the viral life cycle ^29^. Purple Borg has an additional motif composed of GAA, which happens to be the complement of the TC motif (**Figure 2b, Extended Data Table 3**). Based on models for methylation detection we infer that the GAA motif has a 6mA modification in the first A, and we speculate that the YC motifs contain a 4mC modification. This 4mC modification may be introduced by the DNA methylase that is found in all Borgs (**Supplementary Figure 4**). This is supported by the finding that a homolog is also encoded in Sulfolobus virus STSV1 (YP_010357641.1) and STSV2 (YP_007348303.1), and interestingly also in the Haloarcula virus HJTV-2 (YP_010357641.1). From this we conclude that *Methanoperedens* and Borg genomes have clearly distinct methylation patterns, and these could be important in protecting the Borg DNA from degradation by the host machinery, and regulating Borg replication and gene expression.

**Figure 2.**
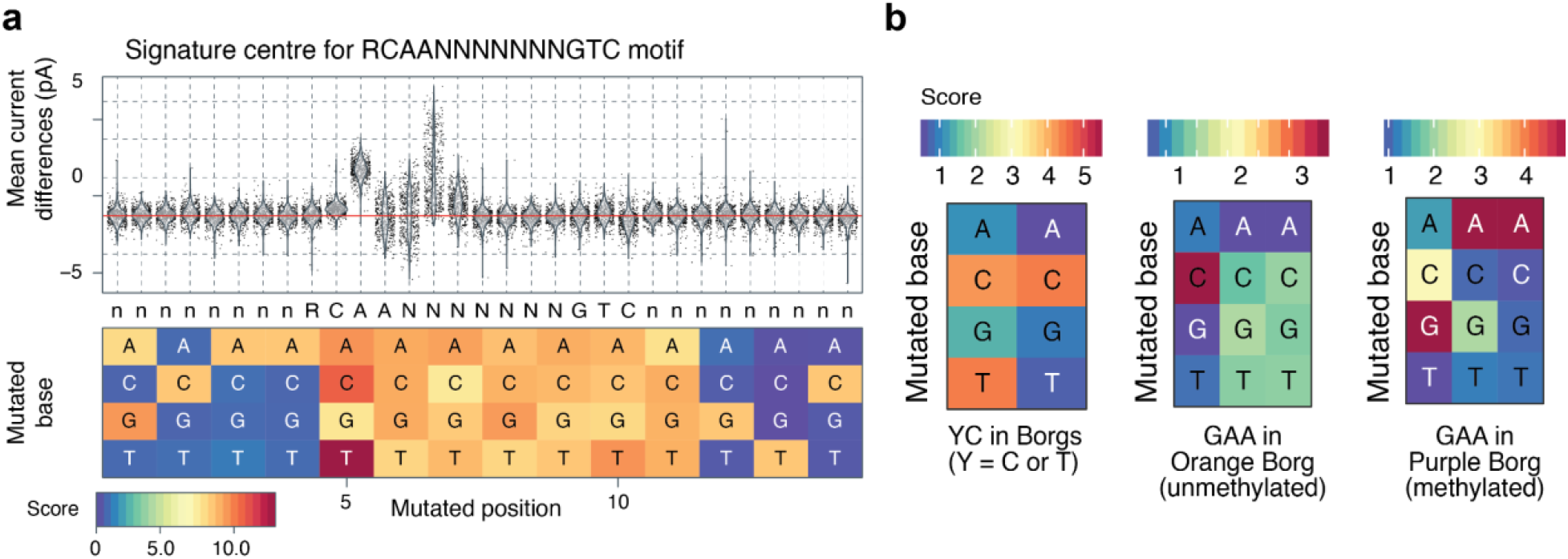
*Methanoperedens* and Borgs have distinct methylation patterns. **a.** Current differences between native and amplified DNA at RCAANNNNNNNGTC motif sites in cMp. **b.** Motif refinement score for related motifs with one substitution.

### Widespread Borg OmcZ nanowire genes that enhance metabolism

Recently, methane-consuming archaea were proposed to use nanowires to transfer metabolic electrons to extracellular electron acceptors, such as minerals or syntrophic partners (extracellular electron transfer, EET) ^30–33^. However the Borg proteins identified are not known to transfer electrons or play a role in metabolism ^33^. Subsequently, it was noted that *Methanoperedens* genomes encode nanowire assembly genes homologous to cytochrome OmcZ, which is processed by OzpA (<u>O</u>mc<u>Z</u> <u>p</u>rotease), and prolyl isomerase, and EET has been confirmed experimentally ^30, 34^. Of 111 cytochromes, OmcZ is the only nanowire-forming outer-surface cytochrome essential for long-range (>10 µm) EET by forming microbial communities ^30, 31^ .

Importantly, we identified potential OmcZ genes in Borgs and evaluated the likelihood that they form nanowires by analyzing conserved structural and assembly features. There is high conservation of the electron-transferring heme region (**Figure 3a**), including the histidine pair that brings hemes closer to confer the 1000-fold higher conductivity of OmcZ compared to other nanowires ^30, 31^ (**Extended Data Figure 7a**).

**Figure 3.**
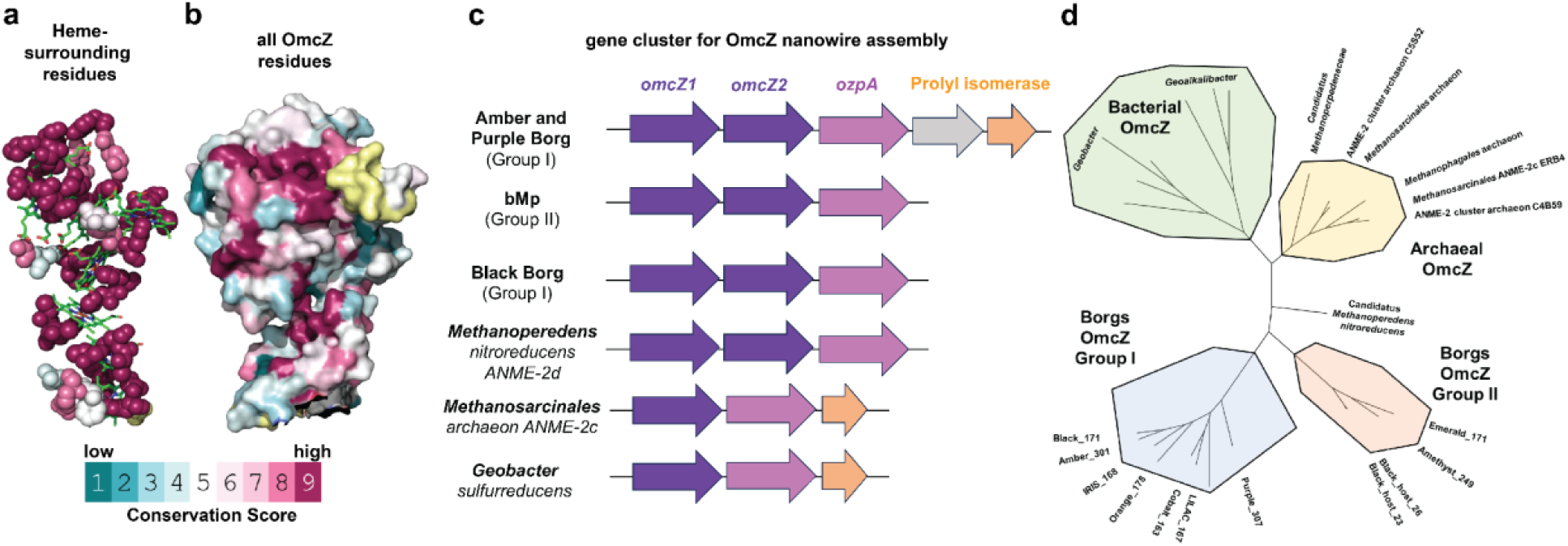
OmcZ nanowire assembly genes that enhance metabolic capacity are widespread in Borgs. Evolutionary conservation of **a.** residues within 3 Å from the heme chain and **b.** surface residues of OmcZ (PDB 7lq5). Yellow regions represent residues lacking enough sequences for comparison. **c.** The nanowire assembly gene cluster is conserved across diverse Borgs and their hosts. **d.** Phylogenetic tree based on OmcZ nanowire assembly proteins.

Both the predicted OmcZ of Borgs and co-occurring *Methanoperedens*s show high conservation of key surface residues that confer high protein stability in denaturing and acidic (pH < 1.6) conditions ^30, 31^ (**Figure 3b**). A subset of Borg OmcZs lacked the non-canonical heme-binding motif (CXnCH, n>2), containing a solvent-exposed loop that traps soluble electron acceptors ^30^. Thus, the host *Methanopereden*s could use these nanowires to pass electrons to insoluble, rather than soluble, electron acceptors. The OmcZ-precursor in Black Borg has two additional β-strand enriched domains compared to bacterial OmcZs (**Extended Data Figure 7b-d**), but the significance of these remains uncertain.

Notably, the homologs of OmcZ form two clades, the first composed only of Borgs (Group I) and the second composed of the *Methanoperedens* hosts and other Borgs (Group II). Both clades are distinct from all other OmcZ homologs (**Figure 3d**). The entire gene cluster required for OmcZ nanowire assembly, comprising OmcZ and the expected peptidase and isomerase, is conserved in several Borgs and in their host’s genomes (**Figure 3c,d, Supplementary Figure 5**). As reconstitution of this cluster in a heterologous host is sufficient to produce OmcZ that self-assembles into nanowires ^30^, our findings suggest that *Methanoperedens* use OmcZ nanowires encoded by Borg genomes to enhance their metabolic capacity to couple methane oxidation to reduction of soil-associated ferric iron oxides or oxyhydroxides.

### Metabolic genes of *Methanoperedens* and Borgs are expressed *in situ*

Reconstructing the metabolism of the genomes of *Methanoperedens* that are predicted Borg hosts confirmed that their main metabolism is likely anaerobic methane oxidation (**Figure 4, Supplementary Table 11**, ^1^). Other than terminal electron acceptor reactions mediated by nanowires, electrons from methane oxidation may be donated to selenate reductase in cMp. Borgs encode a plethora of protein homologs that facilitate an expansion of the host encoded metabolic genes, including CO dehydrogenase, glycolysis and carbohydrate-related genes, transporters involved in the export of lipopolysaccharides, or the import of phosphate and polar amino acids (**Figure 4, Supplementary Table 12**). Both *Methanoperedens* and Borgs also encode numerous S-layer proteins that likely build the outermost layer of the cell. PEGA-domain proteins are additional surface proteins that are exclusively found in Borgs, with four to 21 genes largely colocalized in an individual Borg genome (**Supplementary Table 2, Supplementary Table 12**).

**Figure 4.**
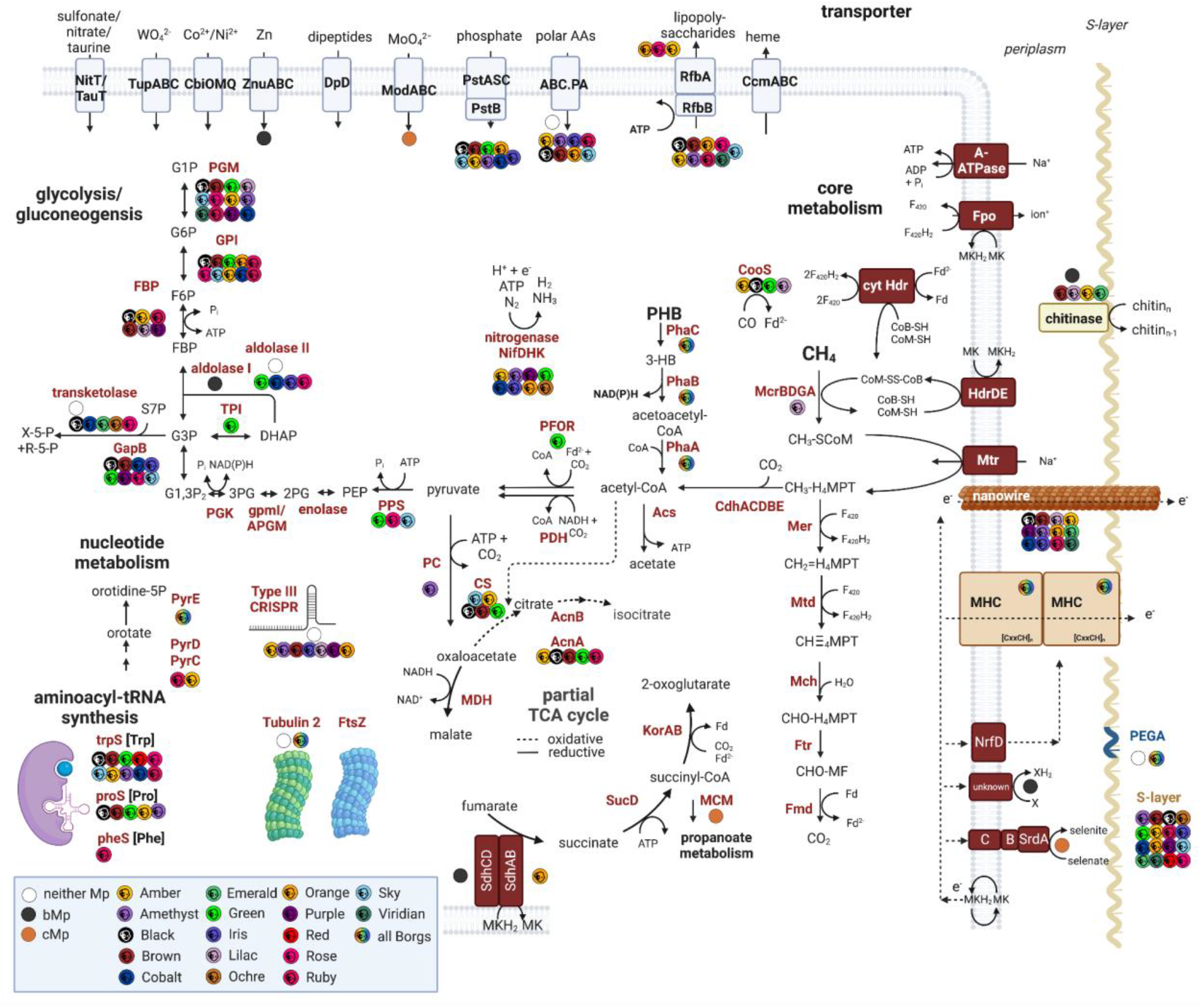
Metabolic reconstruction of two *Methanoperedens* that host Borgs and metabolic potential of Borgs. Elements without white, black, or orange circles are present in both *Methanoperedens* genomes. For an explanation of the abbreviations and locus tags please refer to **Supplementary Table 12**.

We then performed full-length cDNA and gDNA nanopore sequencing on newly collected wetland soil samples (two samples from 50 cm, one sample from 90 cm, 100 cm, 115 cm) in search of transcripts indicating that Borgs and *Methanoperedens* were active at the time of sample collection. Analysis of the metagenomic dataset revealed that these samples consistently contained *Methanoperedens* bMp and Black Borg but no other Borgs. Mapping the cDNA reads revealed that only 8-16% of bMp genes and ≤1% of cMp genes had transcripts detected (**Supplementary Table 13**). The Mcr genes of *Methanoperedens* were most highly expressed (≤225x), followed by genes (≤17x) encoding cell architecture and electron transport-mediating proteins including the S-layer protein, tubulin/FtsZ, adhesion proteins, a phasin, MHCs and nanowire proteins (**Supplementary Table 14**).

Surprisingly, we detected transcripts for 45 – 57% of the genes encoded in Black Borg genome (**Supplementary Table 13**). The nanowire-forming OmcZ genes were among the highest expressed genes of Black Borg. Assuming the inferred Black Borg - bMp linkage is correct, the genomic ratio of Black Borg and its host bMp ranged from 0.4:1 to 0.9:1 in the three samples (**Supplementary Table 15**). Notably, the Borg transcript abundance for the nanowire protein ranged from 3:1 to 6:1 compared to bMp transcripts, indicating that the Borg-encoded nanowire protein is more highly expressed than the host-encoded protein. Other genes in Black Borg with well detected transcripts were other MHCs, a membrane-embedded glycosyltransferase, and core proteins of unknown function (subfam1116 and subfam2007). Overall, this metatranscriptomic analysis illuminated that *Methanoperedens* and Borg genes are expressed simultaneously and that membrane-anchored and extracellular Borg proteins contribute to cell architecture and terminal redox transfer.

### Conserved Borg proteins enable the discovery of new Borgs

The existence of marker proteins distributed over most of the Borg genomes opened a way for improved identification of new Borgs. Previously we used ribosomal protein L11A (rpL11), a near marker Borg protein, to locate and distinguish Borgs. A SingleM search identified one publicly available dataset from 1.5 m depth in a peatland ecosystem warming experiment in Northern Minnesota, USA ^35^ with reads encoding parts of rpL11 from three novel Borgs. Thus, we searched a new assembly of this metagenome with the 40 marker protein sequences and identified 37 to 40 homologs for three Borgs named Maroon (∼65x coverage), Mauve (∼65x coverage) and Liserian (∼17x coverage). The Maroon contigs could be distinguished from those of Mauve as the marker proteins are similar to those of Sky and Rose Borgs whereas Mauve and Liserian proteins are related to those of other Borgs, including Amethyst and Ruby Borgs. Maroon Borg marker protein-bearing contigs, along with candidate contigs with the expected GC content, phylogenetic profile and coverage, were manually curated into a 634 kbp near complete genome with two gaps (originally 11 contigs), ordered based on synteny with the Sky Borg genome. The genome shares all features with previously defined Borgs but displays much fewer TR. Intriguingly, there is no clear indication of *Methanoperedens* in this sample, raising the possibility that Borgs replicate in the methanogenic archaea that are present. Alternatively, *Methanoperedens* may be at very low abundance or the Borgs may exist as environmental DNA, protected from degradation by encapsulation or another mechanism.

### Borg marker gene phylogeny and the “tree of Borgs”

We constructed phylogenetic trees for each of the 40 marker proteins, including homologs from bins for uncurated (or partially curated) Borgs and the 17 complete / near complete Borg genomes. The trees all indicate similar evolutionary relationships amongst the Borgs, including the subdivision of most Borgs into two distinct clades. Given this, we concatenated the 40 sequences of proteins from all Borgs for which we could identify at least 60% of homologs of the marker proteins with reasonably high confidence (in the case of draft genomes). In total, the concatenated marker protein sequences represent 28 different Borgs. The phylogenetic tree generated from the alignment of these sequences confirms the existence of two major clades, one of which was previously represented only by fragments in bins and two complete genomes (**Figure 5**).

**Figure 5.**
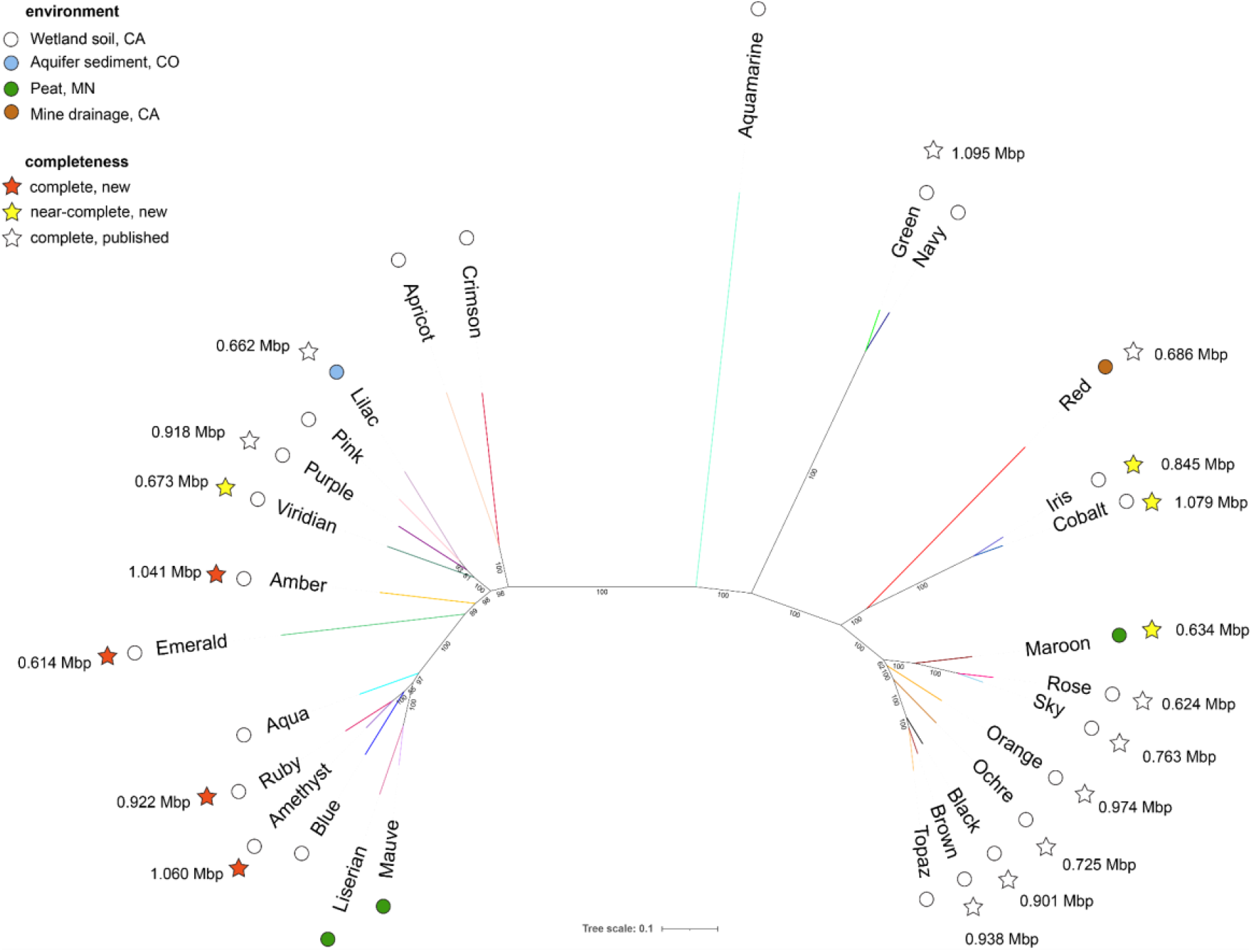
A phylogenetic tree showing the relatedness of 28 Borgs, their genome sizes, completeness and environment of origin.

## Conclusions

This research substantially expands what we know about huge Borg extrachromosomal elements of anaerobic methane-oxidizing *Methanoperedens* archaea. Relative to other ECEs, including plasmids and huge phages, Borgs carry a remarkable inventory of genes for metabolism and for functions related to genome stability and replication. Conserved and generally syntenous Borg single copy marker genes define a shared genomic backbone. Phylogenetic analyses indicate vertical inheritance from a common ancestral type, followed by diversification into two clades. One conserved marker gene encodes an origin of replication binding protein, and other members of the core proteome are predicted to respond to DNA damage and maintain chromosome integrity. We demonstrate that multiheme cytochromes involved in disposing of electrons from methane metabolism are ubiquitous and always present in multiple copies, strengthening the inference that Borgs have evolved mechanisms to augment their host’s ability to oxidize methane to offset the cost of replication of their huge genomes. In fact, metatranscriptomic data reveal that Black Borg nanowires are expressed more highly than those of their putative host *Methanoperedens*.

Although Borgs share many genes, few are ubiquitous, consistent with propensity for gene acquisition via lateral transfer, enabling huge flexibility in gene content. While at first glance resembling plasmids, Borg genomes may encode capsid proteins and have features shared with viruses, including double stranded linear DNA viruses of eukaryotes. The 17 well defined Borg genomes display a consistent genome architecture, further supporting the classification of Borgs as a specific and seemingly novel type of genetic element. Finally, we present the first evidence that distinct methylation patterns may enable *Methanoperedens* to distinguish self vs. Borg genomes. Thus, drawing solely on curated metagenomic sequence information, this study provides a glimpse into the nature and existence of enigmatic genetic elements of some of the most interesting and arguably biogeochemically important archaea yet known.

## Acknowledgements

This publication is based on research in part funded by the Bill & Melinda Gates Foundation (Grant Number: INV-037174 to JFB). The findings and conclusions contained within are those of the authors and do not necessarily reflect positions or policies of the Bill & Melinda Gates Foundation. Funding was also provided by the Innovative Genomics Institute at UC Berkeley (IGI Climate fund, philanthropic donation, to JFB), a DFG fellowship (Project Number: 447383558 to MCS), the NSF-ANR award no. 2210473 (to NSM) as well as the Climate Impact Innovation Fund and the Natural Carbon Solutions Fund (to NSM). SG received support from the Miller Foundation for a visiting professorship at UC Berkeley. BJW is supported by an Australian Research Council Future Fellowship (#FT210100521). CWS is supported by the US Department of Energy (DOE), Office of Biological and Environmental Research, Terrestrial Ecosystem Science Program, under contract DE-AC05-00OR22725 to Oak Ridge National Laboratory/UT-Battelle. We thank Luis Valentin Alvarado and Alex Crits-Christoph for their contribution to field work and generation of sequence datasets, and Shufei Lei and Jordan Hoff for bioinformatics support.

## Author contributions

The study was designed by MCS and JFB. MCS, RS, LDS, BAS and JFB collected samples. DNA and RNA extractions were performed by MCS, LDS, BAS and XD, and whole cells were extracted by MCS and XD. XD and CM performed library preparation and nanopore sequencing. LL generated and analyzed the nanopore datasets with help from CH and JFB. JFB performed binning and carried out the manual genome curation. GC skew, tandem repeat, and genome comparisons were performed by MCS and JFB. Protein family clustering and core Borg proteome analyses were performed by MCS. Structural models were generated by RS and analyzed by MCS and JFB with help from RS. Protein function predictions were inferred by MCS and JFB. JWR performed correlation analysis. Comparisons with viruses were conducted by MCS with input from LDS and RS. DNA modifications in nanopore datasets were analyzed by LL. CS identified and analyzed homologs of gene clusters for OmcZ nanowire assembly under the supervision of NSM. Metabolic reconstruction was performed by MCS, and transcriptome analyses were performed by MCS with help from LL. A search for Borgs in the SPRUCE dataset assembled by BAS was performed by JFB and BW, and the SPRUCE dataset was provided by CWS. NT and SG generated the phylogenetic tree using the sequences dataset provided by MS and JFB. SH, JB, and SJ supervised experiments and data analysis, provided feedback on the study design and methodology, and provided resources and funding.

## Methods

### Sampling and nucleic acid extraction

We recovered deep soil samples from 75 - 120 cm below the soil surface from a wetland in Lake County, California, USA, in September 2021 to construct new metagenomic datasets. DNA was extracted from 5-10 g of soil per sample with the DNeasy PowerMax Soil Kit (Qiagen) and used for short read and long read sequencing. The same site was sampled again at four different depths (twice at 50 cm, once at 90cm, 100cm and 115cm) in November 2022 to construct metatranscriptomic and matching metagenomic datasets. RNA and DNA was co-extracted using a combination of the RNeasy PowerSoil Total RNA Kit and the RNeasy PowerSoil DNA Elution Kit (Qiagen), and then further processed for long read nanopore sequencing.

### Whole cell extraction and DNA crosslinking, and viral concentration

Whole cells were extracted from soil samples by separating particles through low speed centrifugation and subsequent density centrifugation with Optiprep (Sigma). Half of the cell pellet suspension was subjected to formaldehyde treatment (1% v/v) to crosslink protein and nucleic acid within whole cells, and the other half served as a control cell suspension. DNA was extracted and used for long read sequencing. Borg DNA from Black Borg was detected in control cells, but the crosslinked samples did not provide sufficient recovery for sequencing and further analysis. Another attempt to enrich Borgs was undertaken by supersaturating wetland soil with potassium citrate buffer, sequential particle separation through low speed centrifugation three to five times, and passing the supernatant through a 0.22µm membrane and Amicon filter. DNA extracted from the filtrate was sequenced and used in the correlation analysis described in this study.

### Short read and long read sequencing for metagenomics

Paired end 2×250 bp reads were generated from Illumina sequencing on a NovaSeq SP 250PE at the QB3 facility, University of California, Berkeley, USA. Sequencing adapters, PhiX, and other Illumina trace contaminants were removed with BBTools ^36^ (v38.79) and sequence trimming was performed with Sickle ^37^ (v1.33).

Long-read sequencing was performed on three samples using GridION and PromethION P24 sequencing devices at Oxford Nanopore Technologies Inc., New York, USA. Native DNA libraries were prepared with the Ligation Sequencing Kit (LSK114). Amplified DNA libraries were prepared with the Repli-G mini kit (Qiagen) and LSK114. Shorter reads (<1kb) or low quality reads (mean q-score <=10) were filtered out. Adapter sequences were removed using <u>porechop</u> (v0.2.4) ^38^. Branching artifacts caused by multiple displacement amplification were removed by aligning reads to themselves with mappy ^39^(v.2.24), followed by removal of reads with non-diagonal self-alignment. Native and amplified reads were jointly assembled with metaFlye ^40^(v2.9). Contigs were polished with <u>medaka</u> consensus (v1.7.1). Short-read polishing was done with Hapo-G ^41^(v1.3.1).

### Long read sequencing and data processing for metatranscriptomics

Total RNA samples were polyadenylated using *Escherichia coli* poly(A) polymerase (NEB, M0276L) for 5 min and purified by GeneJET RNA Cleanup and Concentration Micro Kit (ThermoFisher, K0841). The poly(A)-tailed total RNA samples were reverse transcribed and PCR-amplified into full length cDNA following the Oxford Nanopore Technologies PCR cDNA Synthesis (PCS109) protocol. cDNA amplicons were barcoded (EXP-NBD114), pooled and sequenced with FLO-PRO114M flowcells on PromethION, and basecalled using Guppy (v6.3.9). Adapter sequences were removed using porechop and reads were trimmed with bbduk (minavgquality=20 qtrim=rl trimq=20). The cDNA reads were then mapped onto the 17 curated Borg genomes and two *Methanoperedens* genomes using minimap2 (-ax map-ont --secondary=no) ^39^(v.2.24-r1122). Alignments were filtered using seqkit bam (--field ReadCov –range-min 70, –field Acc –range-min 95). The alignments per gene were calculated with featureCounts (--fracOverlapFeature 0.1) ^42^ (v2.0.3).

### Manual genome curation

Illumina reads were mapped to nanopore-based assembled sequences classified to be of Borg origin to assess the overall sequence accuracy. Manual curation was undertaken with the goal to generate complete genomes. Obviously chimeric regions, often evident by a large change in GC content and by lack of Illumina read support, as well as unsupported sequence blocks that consisted of mirror images, were removed. Regions lacking perfect read support (0 SNPs allowed) throughout the sequence were flagged and, where possible, errors corrected by mapping reads with lower stringency (e.g., 3% SNPs) to the flagged regions. Individual base pairs in the consensus sequence were then inserted/deleted/replaced manually. Internal segments without read support were removed and replaced by gap filling informed by Illumina reads and using Illumina contigs that were identified by blastn. As Illumina contigs can be chimeric and contain local assembly errors, curation benefited from the availability of Illumina-assembled sequences from multiple samples. The majority Illumina sequence was favored where there was uncertainty, and the newly incorporated sequence verified by stringent Illumina read mapping. Contig ends were extended using unplaced Illumina reads until newly incorporated sequences were sufficient to recruit additional Illumina or nanopore contigs. Assembled sequence ends were extended into terminal ITRs by read mapping until no further extension was possible. Given the generally high read coverage throughout, this was taken to signify the ends of the genome. The genome was then considered complete, so long as there was perfect Illumina read support throughout. In two cases where it was not possible to generate a single Borg sequence extended into ITRs, contigs were ordered and oriented based on alignment with complete Borg genomes. The completeness of genomes not fully extended was assessed based on replichore structure (two replichores of unequal length anticipated), cumulative GC skew and alignment with complete genomes.

### Tandem repeat identification, replication prediction and genome topology verification

Tandem repeats were identified using a custom Python script (https://github.com/rohansachdeva/tandem_repeats) published previously ^5^. Nucleotide TRs were searched using a stringent threshold of ≥50 nt regions and ≥3 TR units and allowing no mismatch (—min_region_len 50—min_repeat_count 3). aaTRs were identified in the concatenated proteins file using a threshold of ≥16 amino acids and ≥3 TRs (-l 3—min_repeat_-count 3—min_uniq_nt 1—min_region_len 16). GC skew (G-C/G+C) and cumulative GC skew were calculated to identify the replication origin and termini, and verify genome topology using the iRep package (gc_skew.py) ^43^.

### Structural modeling and structure-based homology search

Structural modeling of all Orange Borg proteins was performed using AlphaFold2 ^7^ via a LocalColabFold (—use_ptm—use_turbo—num_-relax Top5—max_recycle 3) ^44^. The AF2 rank1 models were profiled against the Protein Data Bank (http://www.rcsb.org/pdb/) (v5e639, date: 2023-04-20) using foldseek ^45^ (v53465) and the easy-search module, or manually queried in PDBeFold ^46^. Protein structures were visualized and superimposed in PyMOL ^47^ (v2.3.4).

### Protein family clustering and functional annotation of protein subfamilies

A dataset of 20,280 Borg proteins was constructed using all 8,322 protein-encoding ORFs of the 7 new Borg genomes, and the 11,958 Borg proteins from the previously published ten complete Borg genomes ^5^. All proteins were clustered into protein subfamilies using MMseqs2 ^48^ (v7e284) in an all-vs.-all search (e-value: 0.001, sensitivity: 7.5, and cover: 0.5). A sequence similarity network was built based on pairwise similarities, and protein subfamilies were defined using the greedy set-cover algorithm. HHblits ^49^(v3.0.3) was used to construct Hidden-Markov models (HMMs) for these protein subfamilies based on the results2msa of MMseqs2. These were then profiled against the PFAM database using HHsearch ^50^ (v3.0.3) and an HMM-HMM comparison.

### Functional annotation of individual proteins

Proteins were profiled using InterProScan ^58^ (v5.51-85.0) and HMMER (hmmer.org) (v3.3, hmmsearch) against the PFAM (--cut_nc) and KOFAM (--cut_nc) HMM databases ^59, 60^. TMHs were predicted with TMHMM ^61^ (v2.0) and cellular localization using PSORT ^62^ (v2.0, archaeal mode). tRNAs were searched with tRNAscan ^57^ (v.2.0.9) and rRNAs with SSU-ALIGN ^63^ (v0.1.1). Protein subfamilies were functionally annotated using HMMER (hmmer.org) (v3.3, hmmsearch) and the PFAM (--cut_nc) HMM database^51^. Homology search was performed with blastp^52^. Metabolic reconstruction was based on the Distilled and Refined Annotation of Metabolism (DRAM) framework ^53^ (v1.2.2) and on custom annotations.

### Correlation analysis

In order to measure correlations between abundance patterns of Borgs and potential *Methanoperedens* hosts, a dataset was compiled consisting of genomic sequences from both Borg and *Methanoperedens* genomes in addition to phylogenetically distinct unbinned contigs containing rpL11 genes profiled as *Methanoperedens* as per Al-Shayeb et al. 2022. These sequences were aligned to 83 metagenomic samples obtained from the wetland soil site. Alignments were performed using bbmap.sh ^36^ using the following parameters: ambiguous=random minid=0.96 idfilter=0.97. For samples sequenced with 150bp reads, the editfilter=5 parameter was used; for those samples sequenced with 250bp reads, editfilter=8 was used. Reads aligning to each genome or rpL11-bearing contig were counted and used to calculate average coverage for each. Coverage values were normalized by sequencing depth by calculating the total number of base pairs sequenced per sample and dividing coverage by this value, then multiplying by a 1e11 scaling factor. Pearson correlations between sequencing depth-normalized coverage patterns for each genome or rpL11 contig across all 83 samples were then calculated using pandas and scipy in python v3.8.2. Clustering and visualization were performed using seaborn, matplotlib, and scipy; code for visualization was generated with the assistance of GPT-4 ^54^.

### Detection of Borg and *Methanoperedens* DNA modification patterns

Methylation motifs and type predictions were found using Nanodisco, which identifies loci with high contrast between the ionic current levels of the native and multiple displacement amplification datasets. Although the Borg motifs were too short to be identified directly, Nanodisco identified several that contained CC, TC, or GAA. The true motifs were narrowed down to YC and GAA by manual inspection of Nanodisco refinement plots, which indicated that additional bases were extraneous.

Independently, methylation was called on individual reads with <u>Megalodon</u>, using Rerio models for all context 6mA and 5mC modifications. However, the YC motif was not found with this method, supporting Nanodisco’s predicted modification type of 4mC. An all-context 4mC Rerio model does not currently exist.

### Genome visualization and alignments

Genomes were visualized in Geneious Prime 2021.2.2 and aligned with the MCM algorithm, or progressiveMauve for aligning multiple contigs. Gene neighborhood analysis was performed with clinker ^55^(v0.0.21).

### Taxonomic classification

16S rRNA gene sequences were used for taxonomic classification of archaeal genomes (Methanoperedens, Methanomicrobiales). The 16S rRNA genes were identified using a custom HMM^56^ (16SfromHMM.py, available at GitHub https://github.com/christophertbrown/bioscripts) and taxonomically classified using the SILVA database ^57^. Taxonomic classification was also routinely performed for all genes as part of an integrated pipeline in ggkbase. The gene annotations are compared to the uniprot and uniref databases and taxonomically classified by the best match in these databases.

### Marker-based identification of a Borg-containing peat metagenome

SingleM (Woodcroft et. al. unpublished, https://github.com/wwood/singlem) was used to identify rpL11 sequences from an in-house database of public metagenomes (published June 2019 or earlier) that was scanned using the SingleM “query” subcommand using --max-divergence 3, using rpL11 OTU sequences derived from Borg reference genomes. This identified the run SRR7028199 as potentially containing Borg genomes. Draft Borg genome bins were identified based on shared low GC, coverage and dominance by novel proteins without phylogenetic affiliations. The bin for the Borg genome that was curated to completion was refined by alignment to the Sky Borg genome and for contigs carrying the Borg marker genes.

### Phylogenetic tree construction

Individual phylogenetic trees were constructed using the protein sequences for the 40 marker proteins to confirm concordant topologies, consistent with vertical inheritance. Blastp searches used the 40 marker proteins to identify homologs in additional Borgs (for which complete and near complete genomes were not available) in the wetland soil and SPRUCE peat data. The single - marker datasets were aligned with MAFFT ^58^ (v7.481) using the option L-INS-I, and alignments were trimmed using BMGE-1.12 ^59^ with the BLOSUM30 substitution matrix. Trimmed alignments were then concatenated into a character supermatrix totalling 9,185 amino acid positions and 28 Borgs. A maximum likelihood tree was then built from this supermatrix with IQ-TREE ^60^ (v1.6.12) using the mixture model LG+C60+F+R4 with ultrafast bootstrap support ^61^ calculated from 1,000 replicates.

### Bioinformatic analysis of Borgs OmcZ homologs

Protein phylogeny tree of nanowire-forming OmcZ was constructed by maximum likelihood method with MEGA X ^62^ and presented by iTOL ^63^. Nanowire-forming OmcZ sequences were extracted from full length OmcZ homologs by removing signal peptide and self-inhibitory part after the subtilisin cleavage site. Signal peptide cleavage sites were predicted by SignalP 5.0 ^64^. The *ozpA* cleavage sites were chosen at the corresponding positions to the OzpA cleavage site in *G. sulfurreducens* OmcZ (after FGNSS) in the multisequence alignment. OmcZ homologs without signal peptides were used for AlphaFold ^7^ modeling. The conservation of OmcZ homologs, which were aligned by Clustal Omega ^65^, was mapped to the structure of nanowire-forming OmcZ (PDB: 7lq5) by Consurf ^66^ and presented by PyMOL ^47^.

## Data availability

Newly released Borg and *Methanoperedens* genomes used in this manuscript are available via: https://ggkbase.berkeley.edu/borgs_mp_nanopore/organisms. The publicly available datasets used in this study are available on NCBI under PRJNA866293. The SPRUCE dataset is accessible under GOLD Project ID# Gp0213362 (https://gold.jgi.doe.gov/search). Protein sequences, structural models and the phylogenetic tree of the 28 Borgs from this study are available through Zenodo (10.5281/zenodo.8162866). Detailed annotations and larger datasets are available in **Supplementary Tables 1-15**.

## Extended Data

**Extended Data Table 1.** Detailed Statistics on available Borg genomes. Newly reported genomes from this study are highlighted in colors. The other 10 genomes were published previously ^5^.

| Borg | bp | status | GC (%) | TR regions | TR regions / 100 kbp | TR region length | TR region (%) | TR unit count | TR intergenic | TR in ORF |
| --- | --- | --- | --- | --- | --- | --- | --- | --- | --- | --- |
| Green | 1094519 | complete | 33.6 | 76 | 6.94 | 10223 | 0.93 | 379 | 46 | 30 |
| Cobalt | 1078846 | curated | 31.8 | 41 | 3.80 | 4552 | 0.42 | 221 | 26 | 15 |
| Amethyst | 1060114 | complete | 34.1 | 65 | 6.13 | 10377 | 0.98 | 429 | 32 | 33 |
| Amber | 1041317 | complete | 32.7 | 45 | 4.32 | 6045 | 0.58 | 259 | 11 | 34 |
| Orange | 974068 | complete | 32.4 | 53 | 5.44 | 5675 | 0.58 | 246 | 33 | 20 |
| Brown | 937932 | complete | 32.2 | 51 | 5.44 | 5913 | 0.63 | 259 | 28 | 23 |
| Ruby | 922385 | curated | 33.5 | 62 | 6.72 | 6967 | 0.76 | 351 | 35 | 27 |
| Purple | 918293 | complete | 32.0 | 66 | 7.19 | 6701 | 0.73 | 326 | 42 | 24 |
| Black | 901883 | complete | 32.2 | 57 | 6.32 | 6128 | 0.68 | 289 | 33 | 24 |
| Iris | 845347 | curated | 31.9 | 35 | 4.14 | 4878 | 0.58 | 215 | 27 | 8 |
| Sky | 763094 | complete | 33.0 | 32 | 4.19 | 3889 | 0.51 | 206 | 21 | 11 |
| Ochre | 725447 | complete | 33.0 | 23 | 3.17 | 2108 | 0.29 | 92 | 12 | 11 |
| Red | 685823 | complete | 32.2 | 23 | 3.35 | 3181 | 0.46 | 173 | 11 | 12 |
| Viridian | 673463 | curated | 34.3 | 28 | 4.16 | 3110 | 0.46 | 152 | 15 | 13 |
| Lilac | 661708 | complete | 32.0 | 61 | 9.22 | 6733 | 1.02 | 299 | 33 | 28 |
| Rose | 623782 | complete | 32.8 | 20 | 3.21 | 2132 | 0.34 | 98 | 8 | 12 |
| Emerald | 613519 | curated | 34.0 | 19 | 3.10 | 3955 | 0.64 | 88 | 1 | 18 |

**Extended Data Figure 1.**
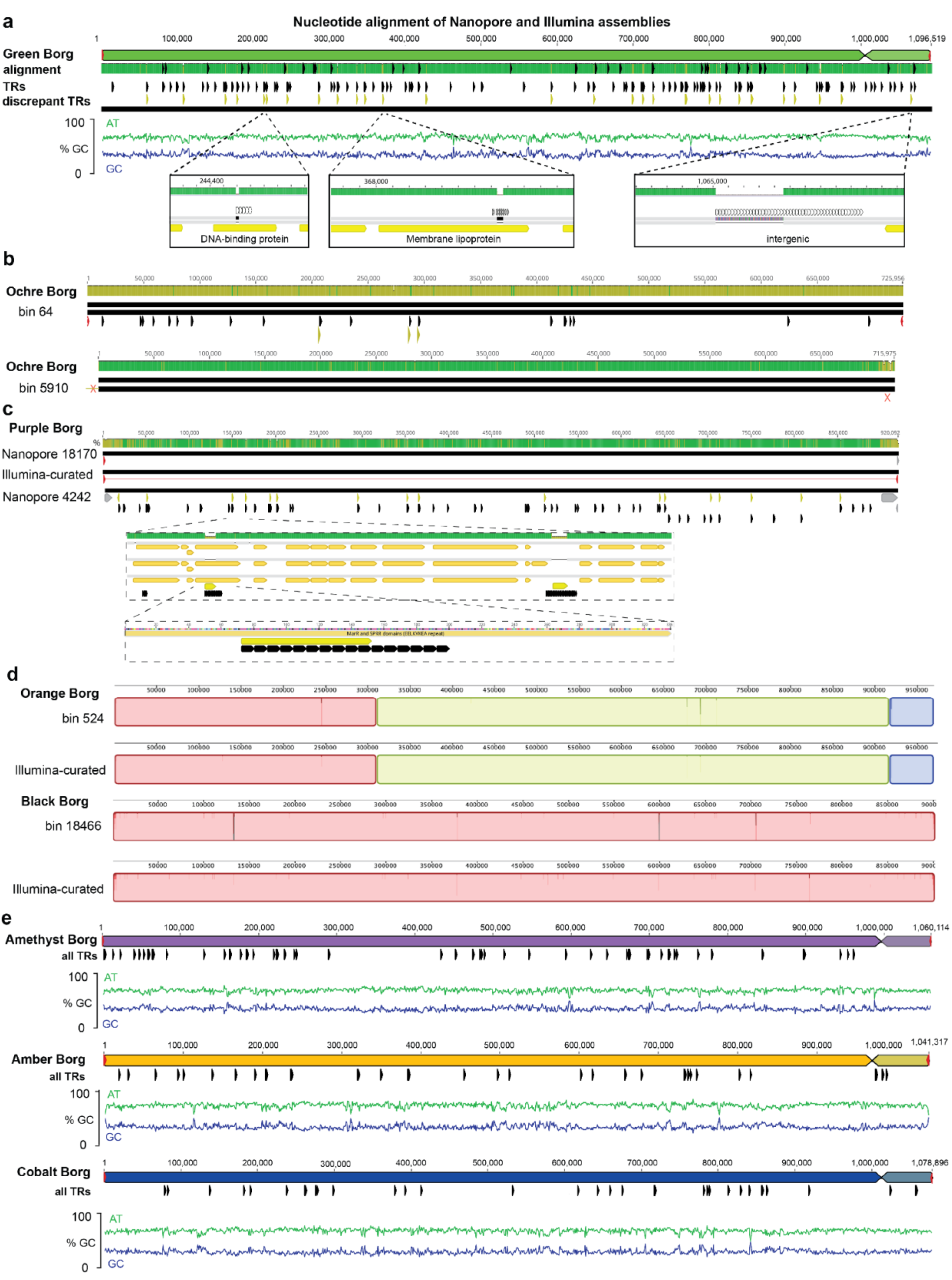
Confirmation of Borg genome architecture including characteristic features reported based on Illumina assemblies, and overviews of new Borg genomes. **a.** Alignment of nanopore and Illumina assemblies for Green Borg. The nanopore Green Borg genome differs from the manually curated Illumina genome almost exclusively in the number of unit repeats in the TR regions (discrepant TRs shown as yellow marks). All TR regions are shown in black, expanded TR regions come to light in the nanopore genome and are shown as discrepant TRs in yellow. **b**. Comparison of the published Ochre genome manually curated from Illumina reads to two versions based on assembly of nanopore reads. The first relatively low accuracy nanopore sequence (bin 64) confirms the overall topology of the Illumina-based genome, including the presence of inverted terminal repeats. The second nanopore sequence has high base accuracy but contained a ∼10 kbp chimeric start (much higher GC content, trimmed from the 5910 sequence used in the alignment) and had a low accuracy terminal region (marked X). **c**. The Illumina-based Purple Borg genome aligned to two nanopore-derived sequences providing overall validation. Also shown are examples of a genic and intergenic regions where nanopore unit repeat count differs. **d**. The overall topology of genomes for Orange and Black Borgs, including terminal inverted repeats, were confirmed using nanopore-assembled sequences; differences were again localized in TR regions. **e.** Overview of three new curated complete genomes recovered from nanopore assemblies. The linear genomes are composed of two replichores and terminated by inverted terminal repeats shown in red.

**Extended Data Figure 2.**
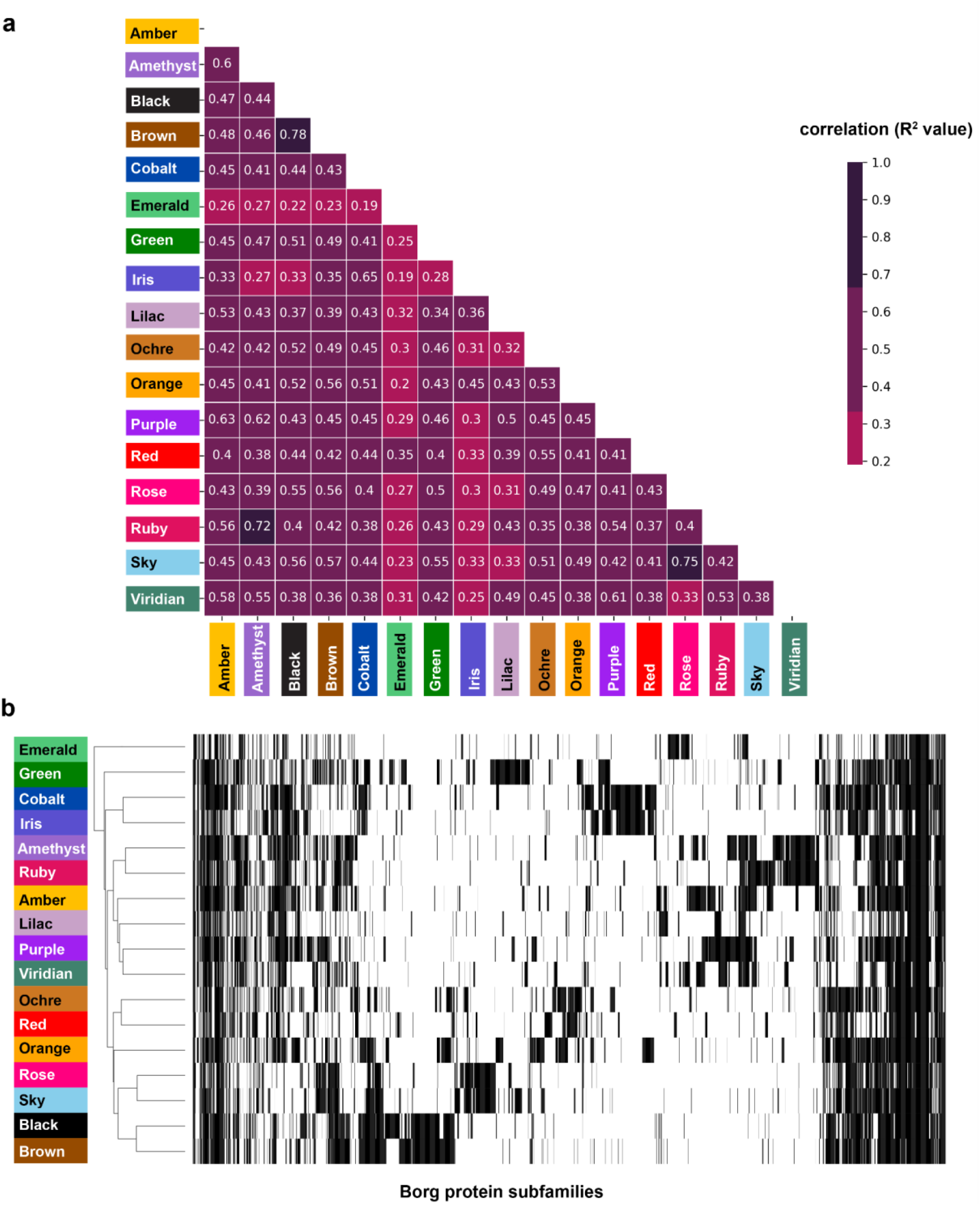
Borg proteome content. **a**. Shared protein subfamily correlation analysis reveals proteome-based relatedness of the 17 curated complete and near-complete Borgs. **b**. Heatmap showing presence (black) or absence (white) of protein subfamily members in the 17 Borgs. Borgs sharing more protein subfamilies cluster together (i.e., Black and Brown Borg).

**Extended Data Figure 3.**
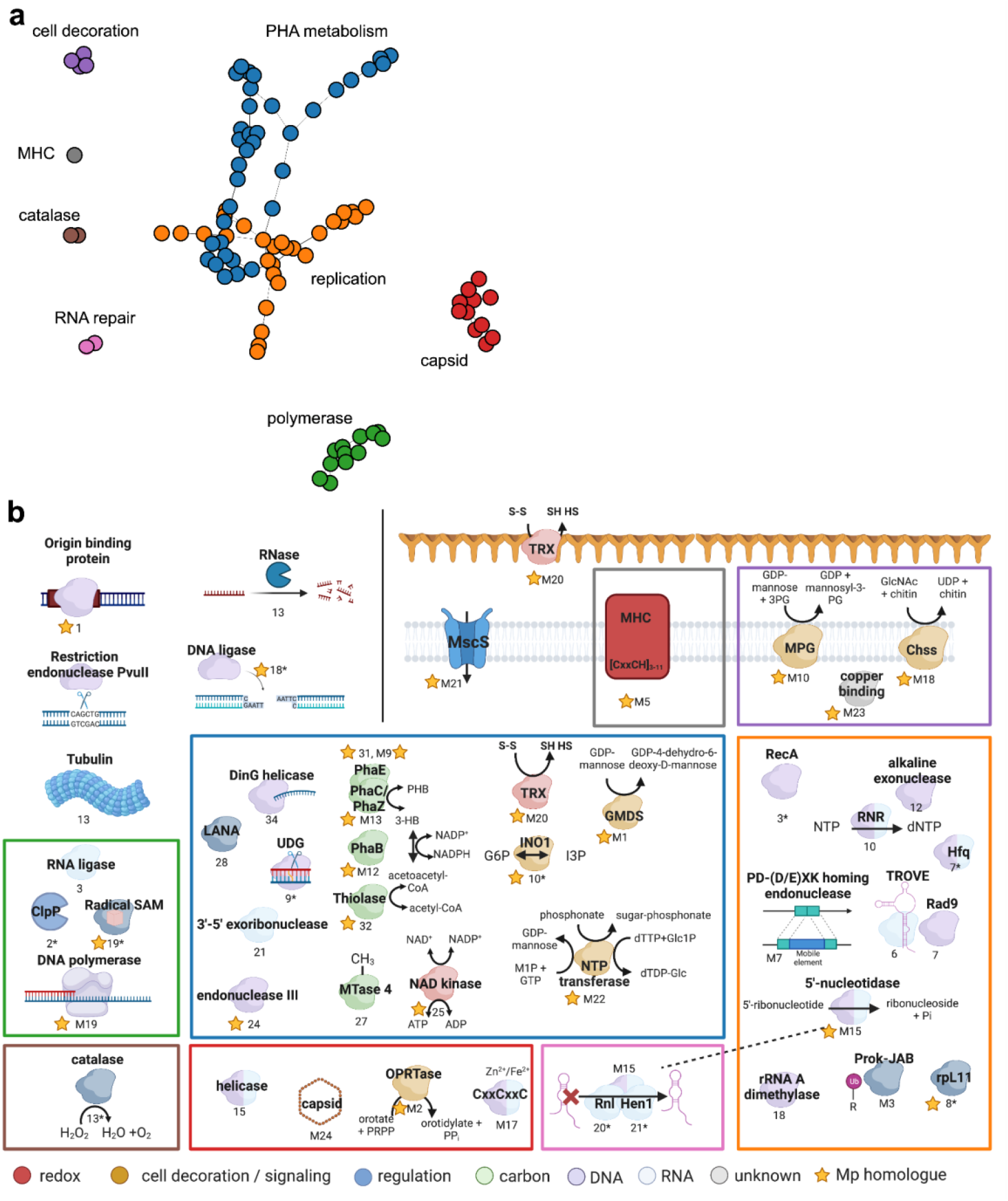
Protein network of core Borg genes and their putative functions. **a**. Core Borg proteins were plotted in distinct clusters based on their consistent colocalization ten genes up- or downstream of each other (n > 5). **b**. Core Borg proteins with functional annotations are shown. Proteins of the same cluster are grouped together in colored boxes. Numbers below objects indicate that they are either marker proteins (numbers), near marker proteins (number and *), or multicopy proteins (number preceded by the letter M). More detailed descriptions of the objects are listed in **Supplementary Table 3**.

**Extended Data Figure 4.**
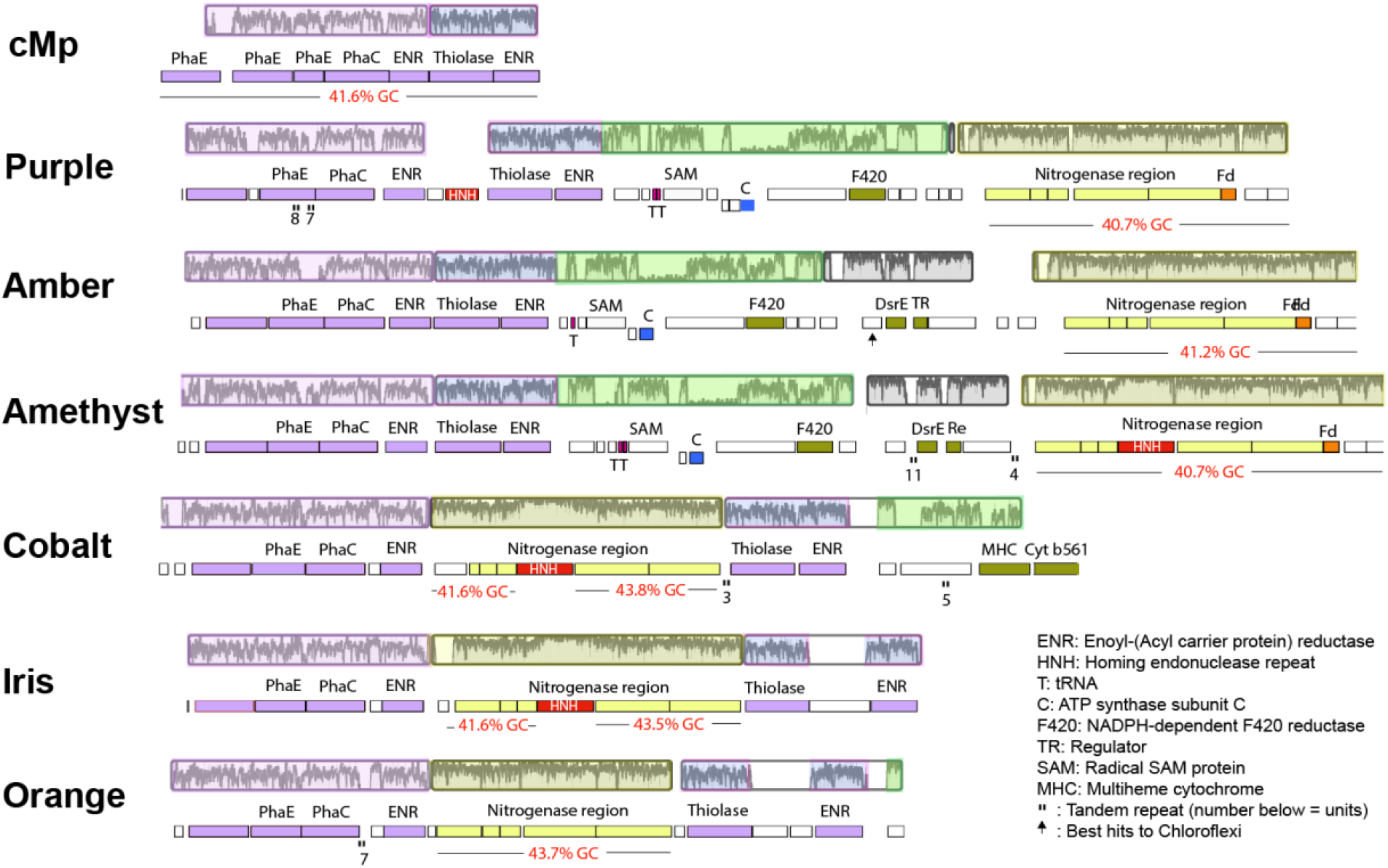
A region encoding genes involved in N_2_ fixation has been inserted within the region encoding for PHA metabolism (purple and blue highlights) in the Cobalt, Iris, and Orange Borg genomes. This nitrogenase-related region occurs downstream in three other Borg genomes. The Borg nitrogenase-related genes have elevated GC contents that match that of the complete *Methanoperedens* genome. Graphs indicating nucleotide-level sequence identity across the Borg and *Methanoperedens* genomes are shown above the coding strand and gene annotation information; highlight colors indicate the largely syntenous blocks.

**Extended Data Figure 5.**
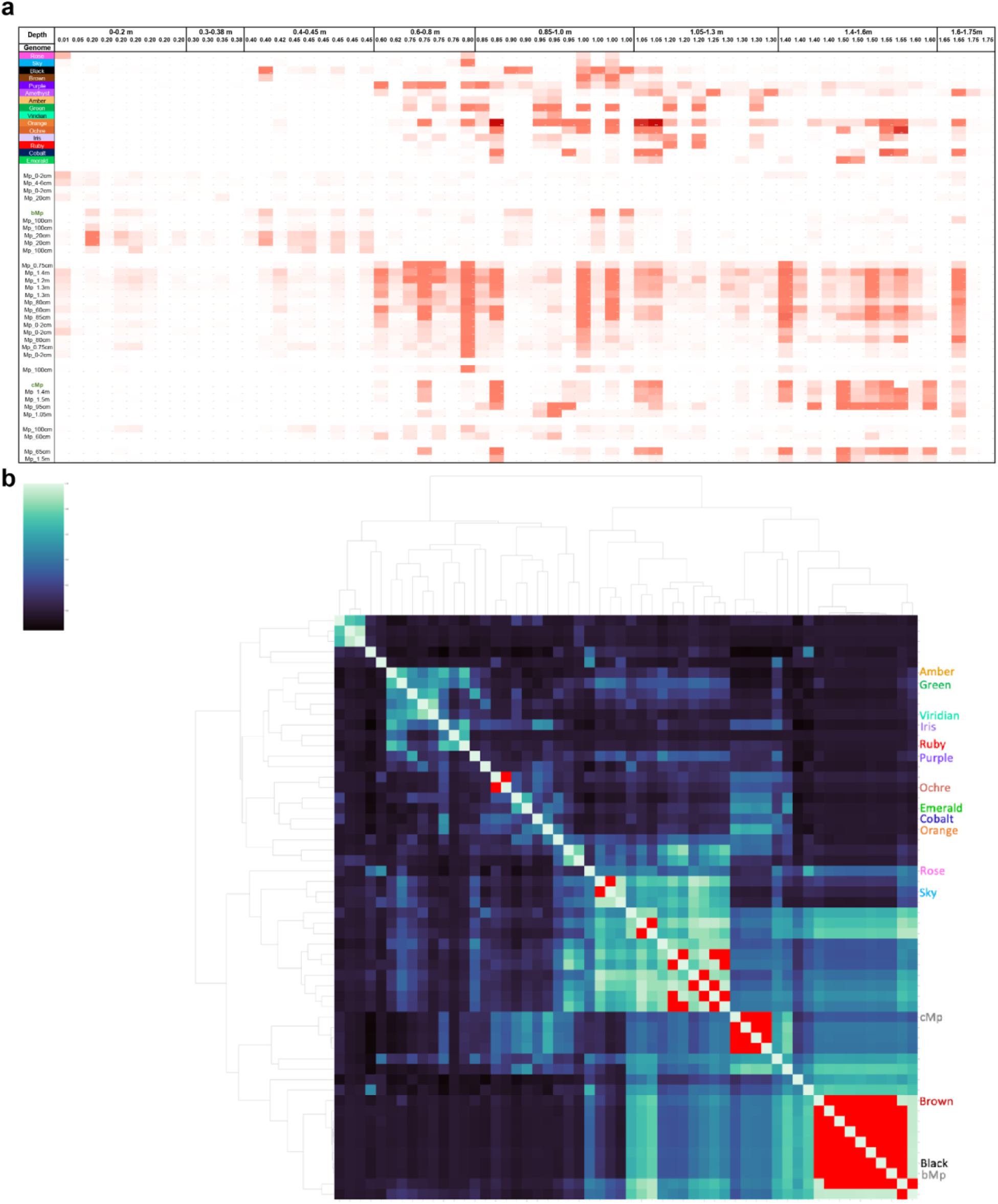
Borg and *Methanoperedens* species abundances over samples, normalized for sequencing depth. **a.** Normalized abundance over samples and depth**. b.** Correlation analysis between coverage patterns of *Methanoperedens* genomes, population representative *Methanoperedens* rpL11 contigs, and Borg genomes across 83 samples. Red cells indicate correlation scores above 0.95.

**Extended Data Figure 6.**
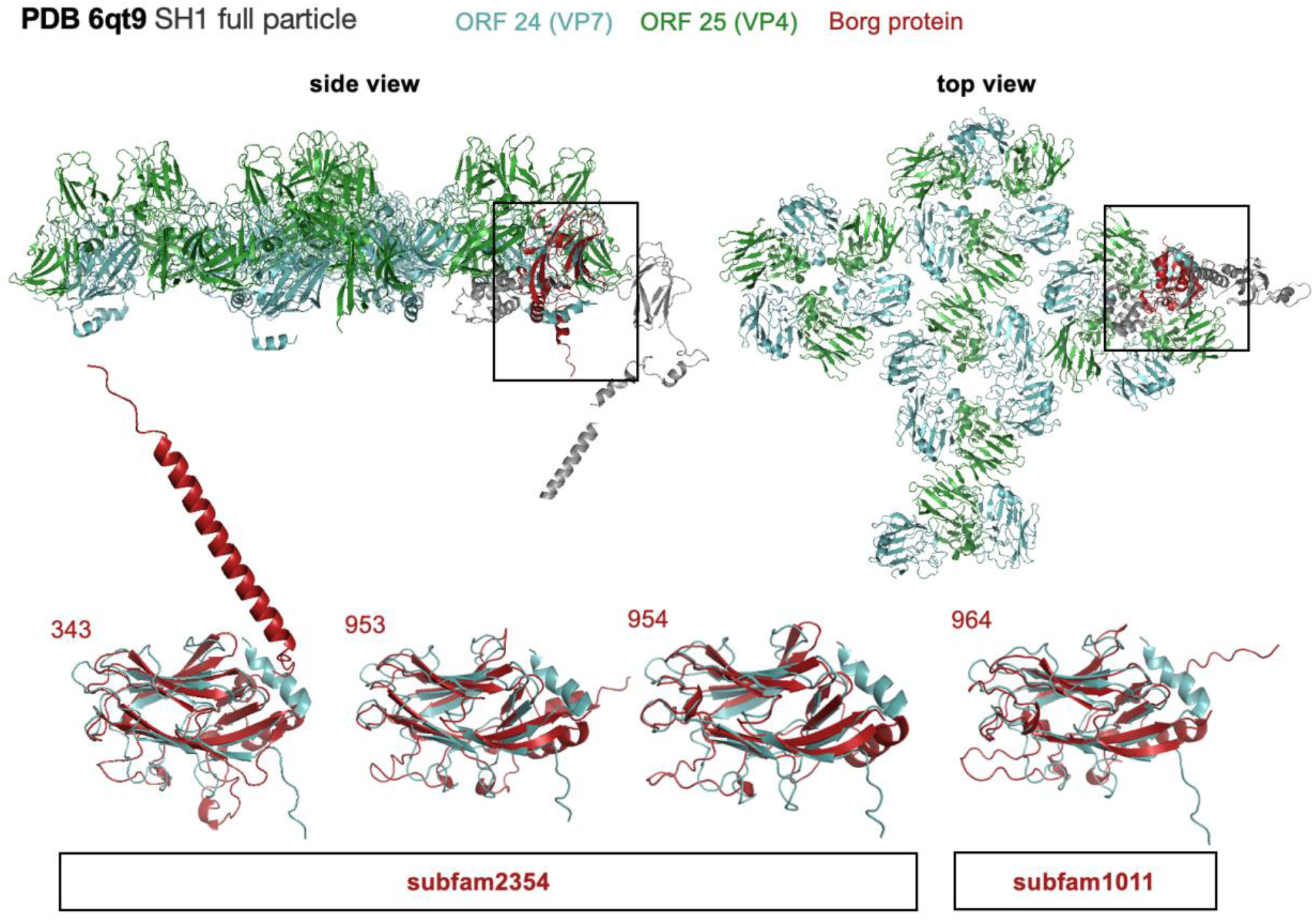
Structural model of putative capsid proteins from Orange Borg and superimposed on the structural matches from an archaeal virus. Structures were predicted with AF2, best structural matches were identified in the PDB using foldseek and visualized and superimposed in PyMOL. Orange Borg proteins and their corresponding locus tags/gene numbers are shown in red. The major capsid proteins of the *Haloacrcula hispanica* virus SH1 are ORF 24 (cyan) and ORF 25 (green), minor capsid proteins are shown in gray. The SH1 full particle structure was downloaded from PDB (6qt9).

**Extended Data Figure 7.**
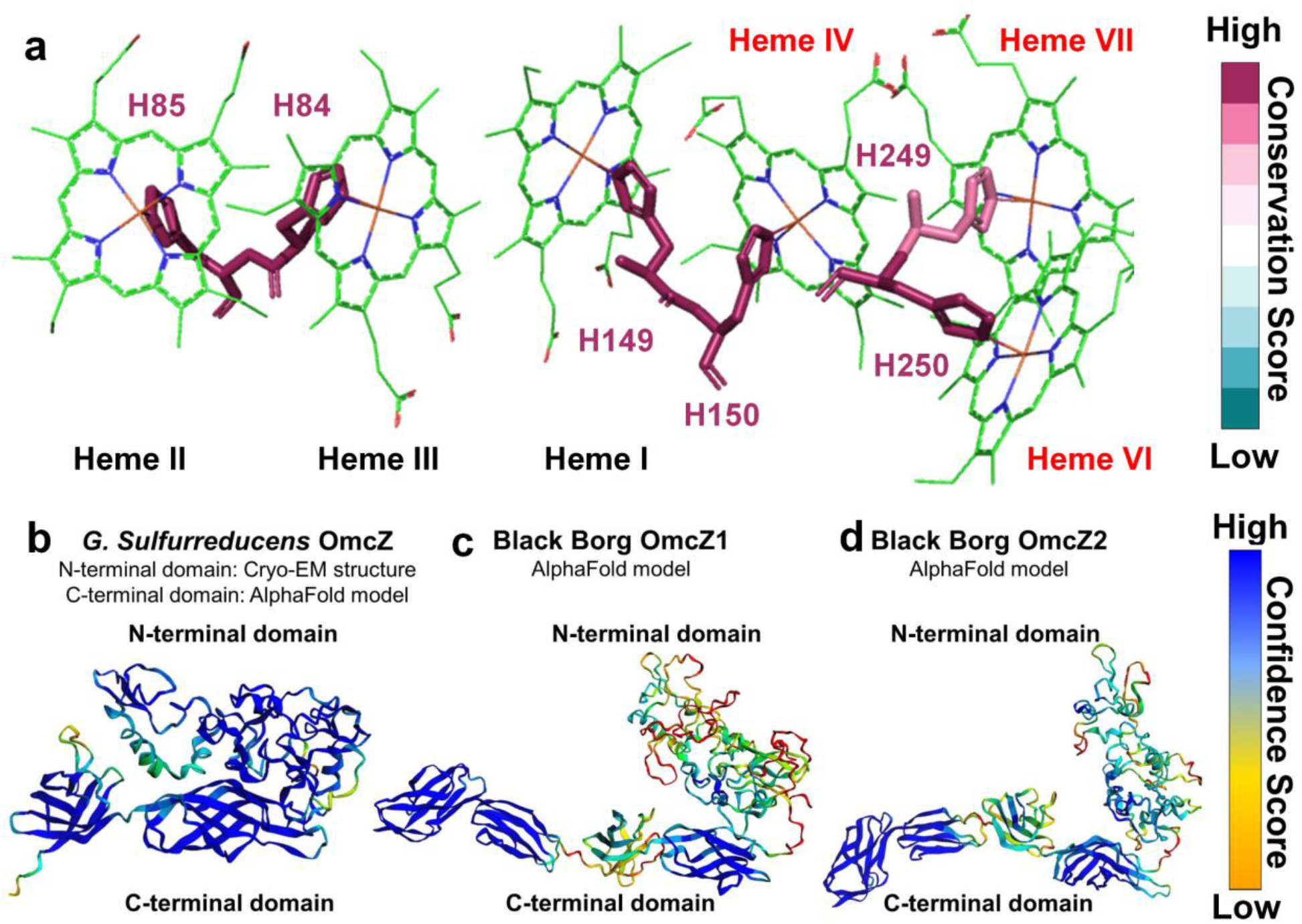
Key structural features important for electron transfer in OmcZ nanowires show high evolutionary conservation in Borgs and hosts. **a.** Histidine pairs that lock hemes more tightly to confer high nanowire conductivity are highly conserved in Borgs and hosts. The residue numbering follows the amino acid sequence of Black Borg OmcZ1 and includes the signal peptide. **b.** AlphaFold model predicts with high confidence that bacterial OmcZ-precursor shows only two β-strand enriched domains in the C-part whereas **c.** OmcZ1- and **d.** OmcZ2-precursors from Black Borgs show four β-strand-enriched domains in the C-part.

**Extended Data Table 2.** DNA replication machinery of Herpesvirus and putative functional homologs in Borgs. The table is based on Weller & Coen, 2012, and was populated with Borg proteins from this study.

| Gene | Abbreviation | Major function | Essential for DNA synthesis in cultured cells | PFAM | Borg subfam with potentially similar function | core protein | PFAM annotation |
| --- | --- | --- | --- | --- | --- | --- | --- |
| UL9 | OBP | Origin-binding protein, helicase | Yes | Herpes_ori_bp | subfam0615 | **marker protein** | Herpes_ori_bp |
|  |  |  |  |  | subfam0300 | no, Green and Ruby | Flavi_DEAD \| Herpes_ori_bp |
| UL30 | Pol | Catalytic subunit of DNA polymerase | Yes | (DNA_pol_B), (DNA_pol_B_exo1) | subfam0942 | **marker protein** | DNA_pol_B \| DNA_pol_B_exo1 |
| UL42 |  | Processivity subunit of DNA polymerase | Yes |  | subfam0273 | **marker protein** | Rad9 |
| UL5 |  | Subunit of helicase/primase; contains helicase motifs | Yes | (Herpes_Helicase) | subfam2200 | **marker protein** | Helicase_C_2 |
| UL8 |  | Subunit of helicase/primase; interacts with other proteins | Yes | (Herpes_HEPA) |  |  |  |
| UL52 |  | Subunit of helicase/primase; contains primase motifs | Yes | (Herpes_UL52) |  |  |  |
| UL29 | ICP8 | Single-strand DNA-binding protein | Yes |  |  |  |  |
| UL23 | TK | Thymidine kinase | No | (Herpes_TK) |  |  |  |
| UL39 | RR1, ICP6 | ICP6 Large subunit of ribonucleotide reductase | No | (Ribonuc_red_lgN) | subfam02016 | **marker protein** | Anaerobic ribonucleoside-triphosphate reductase |
| UL40 | RR2 | Small subunit of ribonucleotide reductase | No | (Ribonuc_red_sm) |  |  |  |
| UL2 | UNG | Uracil-DNA glycosylase | No | (UDG) | subfam2424 | **marker protein** | Uracil DNA glycosylase superfamily |
| UL50 | dUTPase | Deoxyuridine triphosphatase | No | (dUTPase) | subfam2570 | no, Viridian and Emerald | dUTPase |
| UL12 | 5'–3'Exo<br>Viral recombinase | Alkaline nuclease; interacts with ICP8 | No | (Herpes_alk_exo) | subfam2517 | **marker protein + 2** | YqaJ-like viral recombinase domain \| Viral_alk_exo |
|  | dNK | auxiliary |  | (dNK) | subfam1286 | no, Brown and Cobalt | dNK Herpes_TK |
|  |  |  |  |  | subfam2469 | no, Green and Amethyst | KTI12 dNK |

**Extended Data Table 3.** Methylation motif and frequency in the cMp genome and Borgs.

| genome | Motif | Predicted Type | Count Total | Intergenic % | In ORF % | Motifs per 100kbp |
| --- | --- | --- | --- | --- | --- | --- |
| **cMp** | AWGCCC | 4mC | 3773 | 60 | 40 | 94 |
|  | CCWGG | 5mC | 15370 | 53 | 47 | 384 |
|  | CGANNNNNNGGC | 6mA/5mC | 795 | 43 | 57 | 20 |
|  | GCCNNNNNNTCG | 4mC | 795 | 65 | 35 | 20 |
|  | GGGGGA | 6mA | 1190 | 32 | 68 | 30 |
|  | GGNCC | 4mC | 8226 | 53 | 47 | 205 |
|  | RCAANNNNNNNGTC | 6mA | 674 | 58 | 42 | 17 |
|  | RCAGNNNNNNNTCC | 6mA | 934 | 54 | 46 | 23 |
| **Black** | GAA | 6mA | 42331 | 40 | 60 | 4694 |
| **Ochre** | GAA | 6mA | 33747 | 35 | 65 | 4652 |
| **Orange** | GAA | 6mA | 45915 | 38 | 62 | 4714 |
| **Purple** | GAA | 6mA | 41842 | 38 | 62 | 4556 |
| **Black** | YC | 4mC | 168573 | 72 | 28 | 18691 |
| **Ochre** | YC | 4mC | 139525 | 73 | 27 | 19233 |
| **Orange** | YC | 4mC | 185445 | 72 | 28 | 19038 |
| **Purple** | YC | 4mC | 169048 | 72 | 28 | 18409 |

## Supplementary Information

**Supplementary Figure 1.**
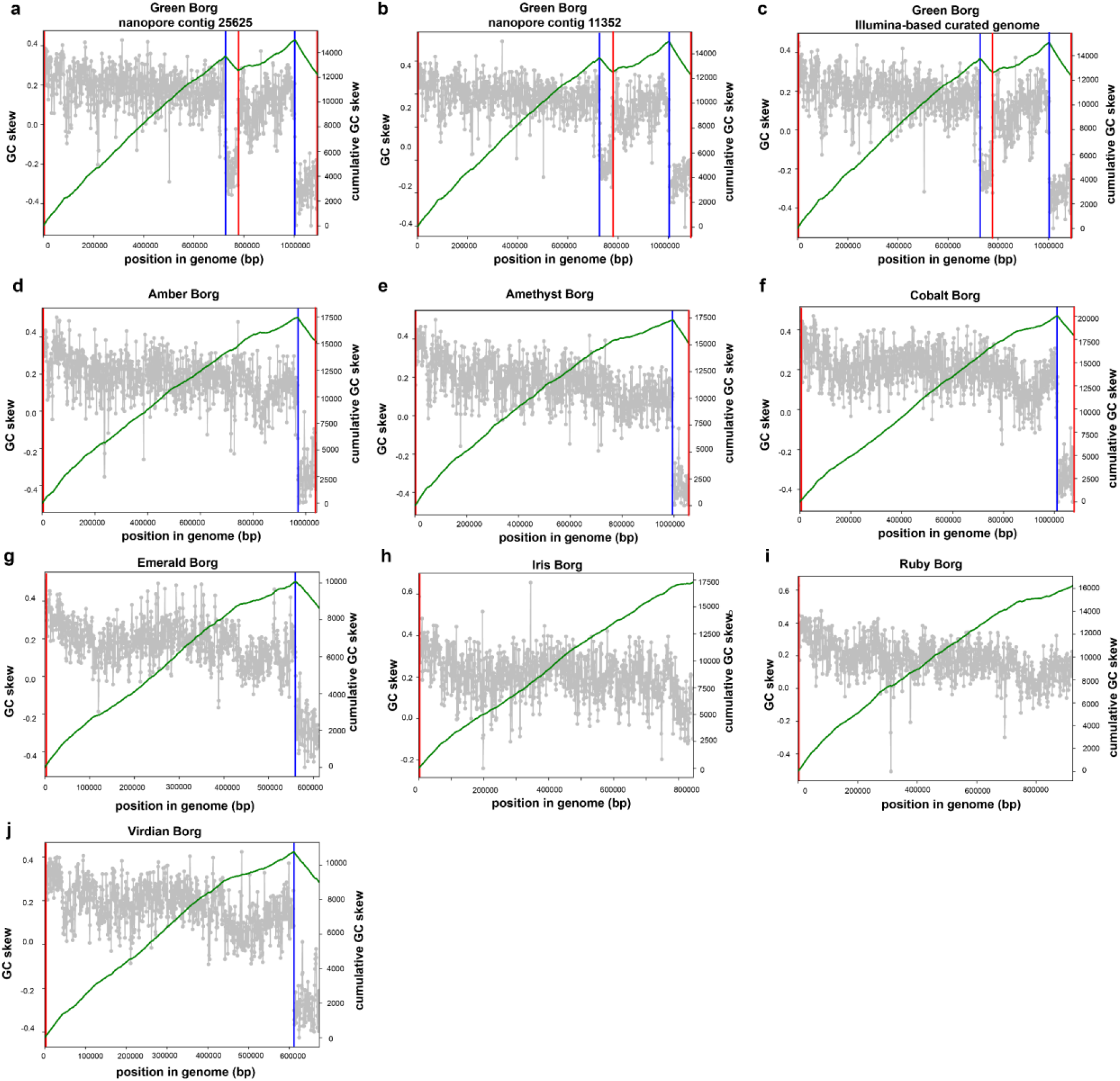
GC skew analysis of nanopore-derived Green Borg contigs and Illumina-based curated genome, and GC skew of new Borg genomes. Green Borg contigs derived from nanopore assemblies were recovered in two different soil samples and the GC skew plots confirm the overall consistent topology (A, B) that was established based on manually curated Illumina assemblies only (C, already published in ^5^. Recovery of 7 new Borg genomes from the nanopore assemblies shows a consistent (cumulative) GC skew for these genomes, suggesting replication from the termini. Iris, Ruby and Viridian Borg are not complete, as reflected by the missing drop in the GC skew peak in the cumulative GC skew, which is absent due to the missing small replichore.

**Supplementary Figure 2.**
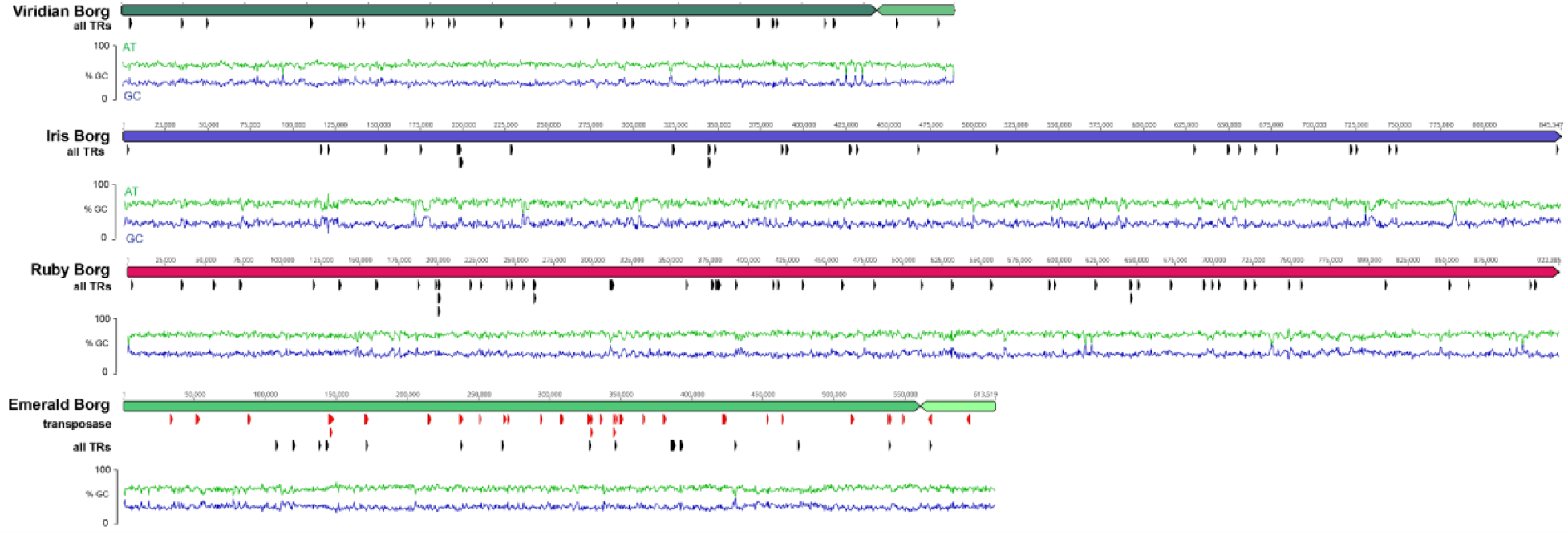
Overviews of the non-complete but manually curated Borg genomes. Red markings indicate the locations of numerous transposase genes in the Emerald genome.

**Supplementary Figure 3.**
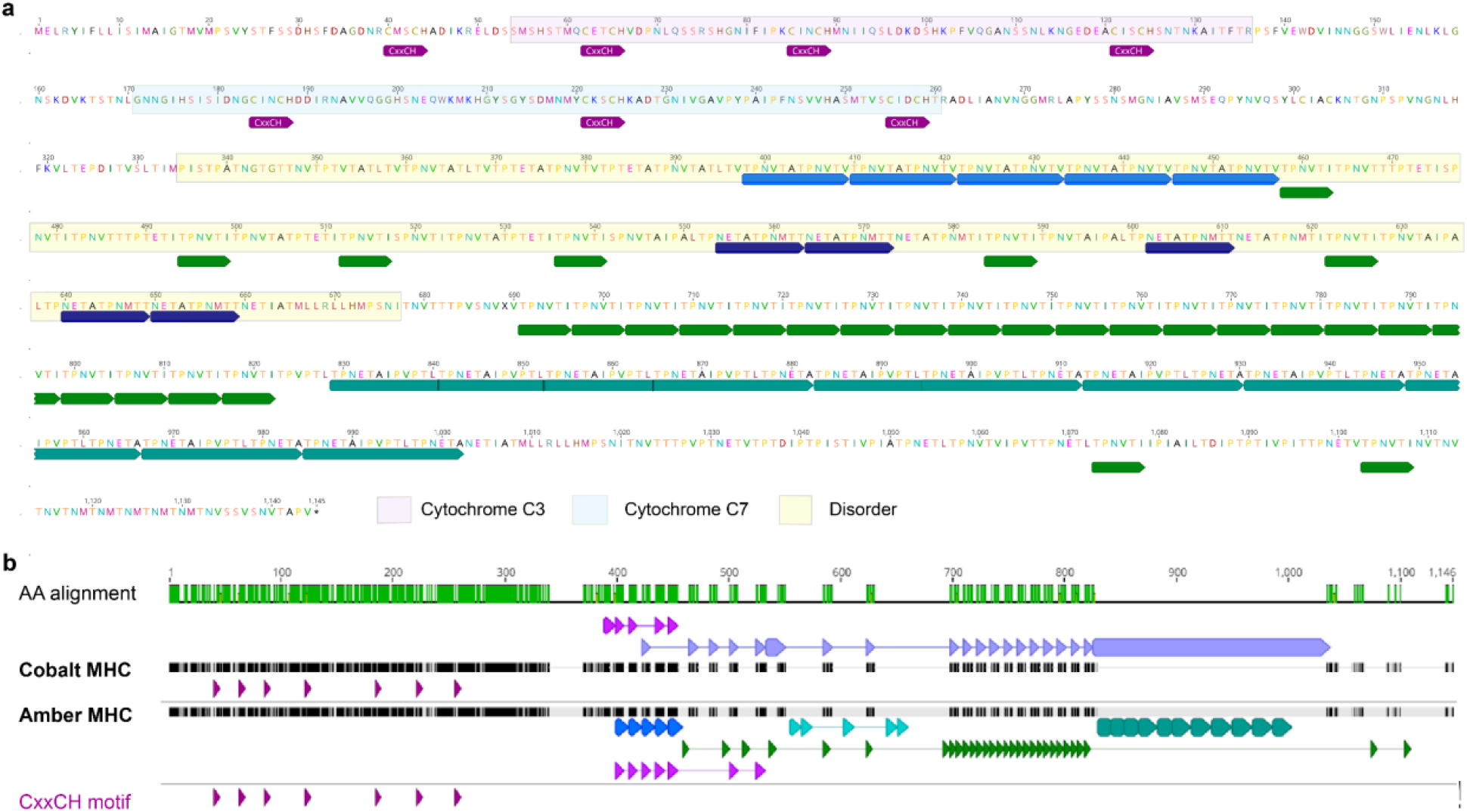
Repetitive regions in Borg DNA. **a**. Amber Borg genome encodes a multi-heme cytochrome (7 heme-binding motifs: CxxCH) that is a complex repeat-bearing protein. Colored arrows with the same color indicate the same repeat sequence. **b**. Alignment of the Amber MHC with TR with that of the Amethyst MHC with TR. Note that a portion of some unit repeats in the two proteins are shared.

**Supplementary Table 1.** Taxonomic classification of all Borg proteins based on uniprot and uniref comparisons.

| domain | count | % classified | % overall |
| --- | --- | --- | --- |
| Eukaryota | 110 | 1.5 | 0.5 |
| unknown | 774 | 10.2 | 3.8 |
| Archaea | 4007 | 53.0 | 19.6 |
| Bacteria | 2472 | 32.7 | 12.1 |
| Fungi | 43 | 0.6 | 0.2 |
| Viruses | 54 | 0.7 | 0.3 |
| Metazoa | 78 | 1.0 | 0.4 |
| Viridiplantae | 16 | 0.2 | 0.1 |
| all genes | 20469 |  |  |
| total unclassified | 12875 |  |  |
| total classified | 7554 |  |  |

**Supplementary Table 6.**
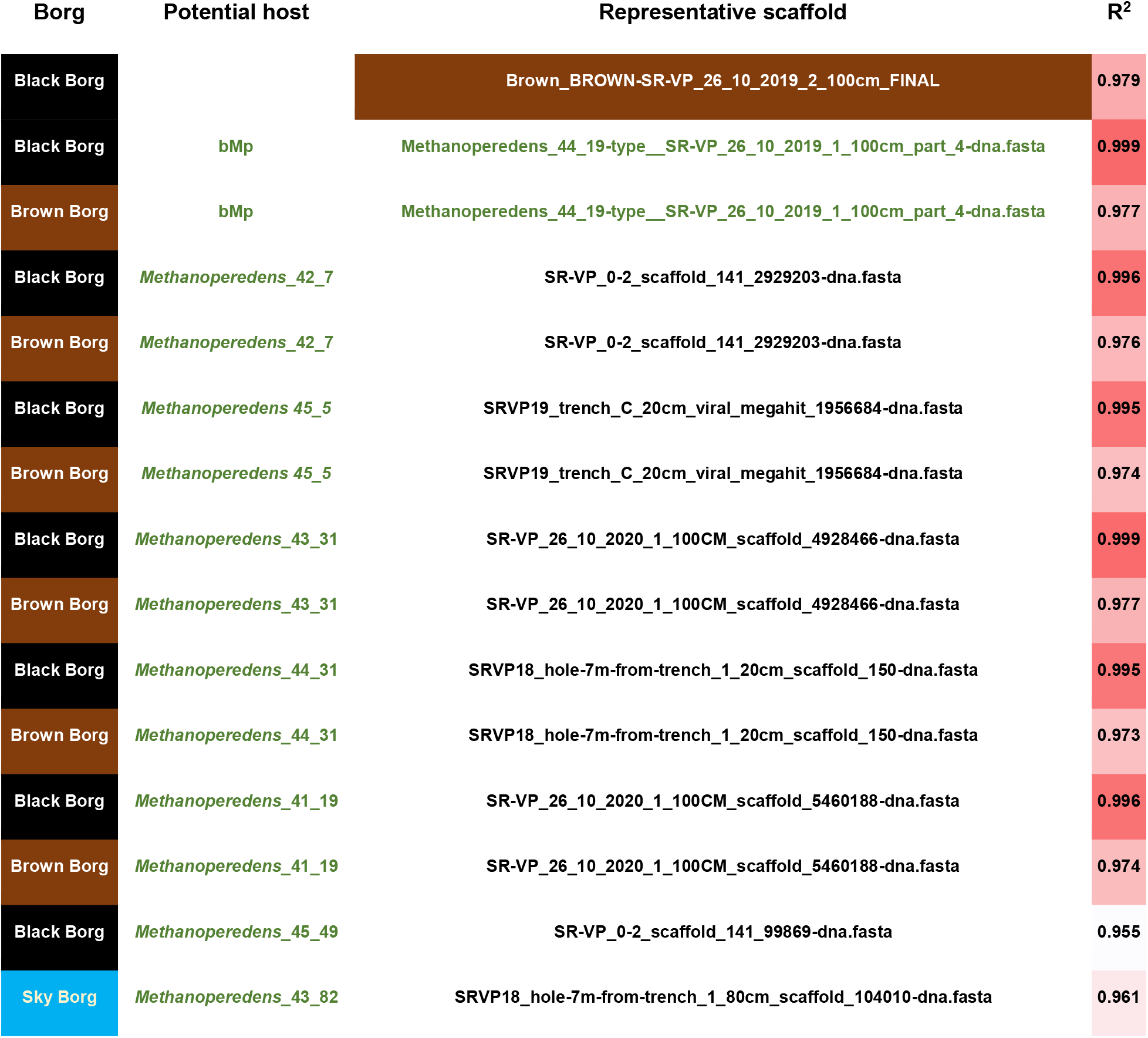
Statistically significant correlations in the pattern of abundance over samples for Borgs. Coverage values for Borg genomes, *Methanoperedens* scaffolds carrying rpL11, the cMp and bMp genomes and draft *Methanoperedens* genomes across 65 soil samples were used to detect possible Borg-host linkages. The data support the prior inference that Black Borg replicates in the bMp genome and suggest that Brown Borg may also replicate in this or a related species.

**Supplementary Table 8.**
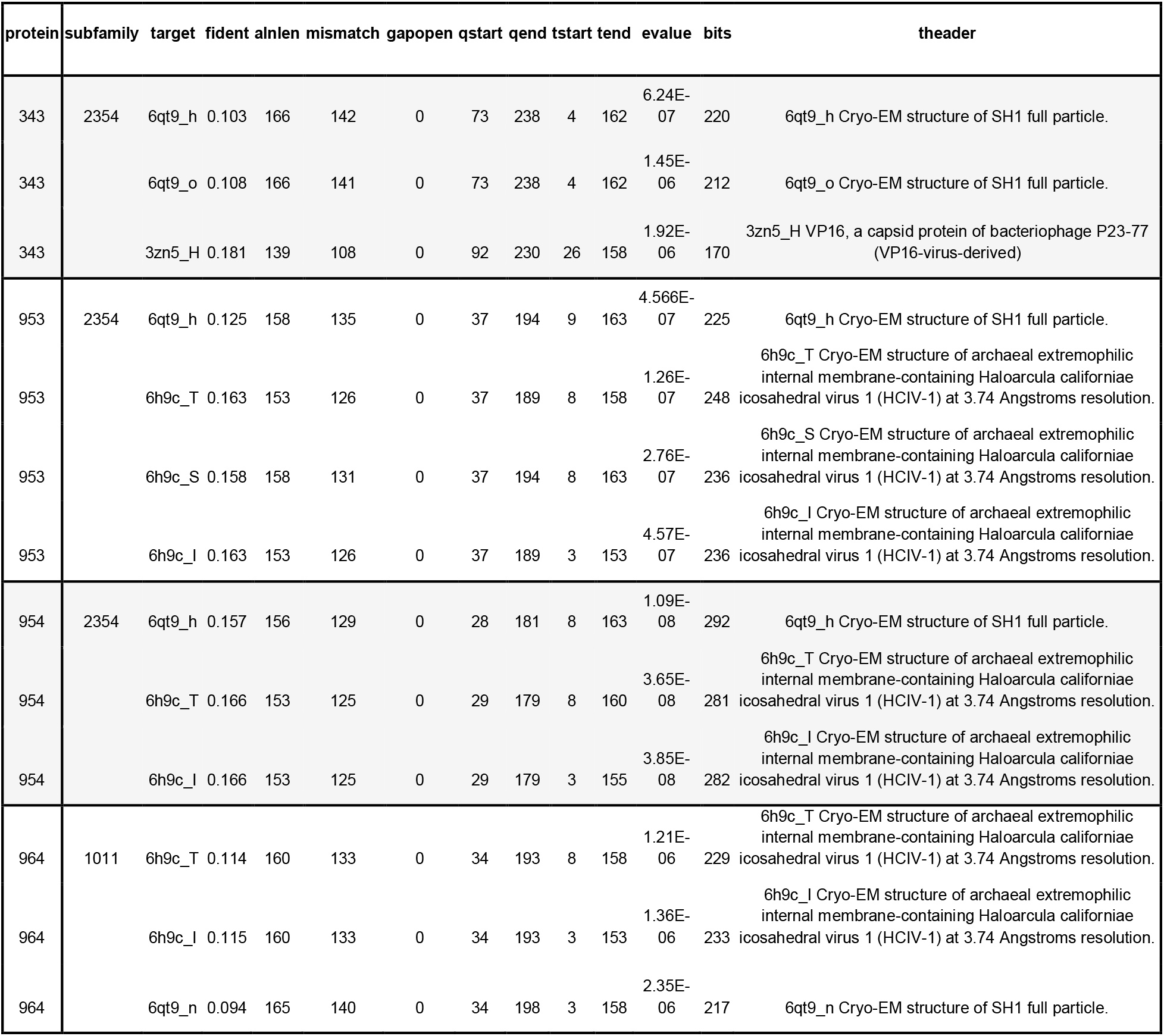
Capsid-related structural matches of Orange Borg proteins from AF2-modeled protein structures.

**Supplementary Figure 4.**
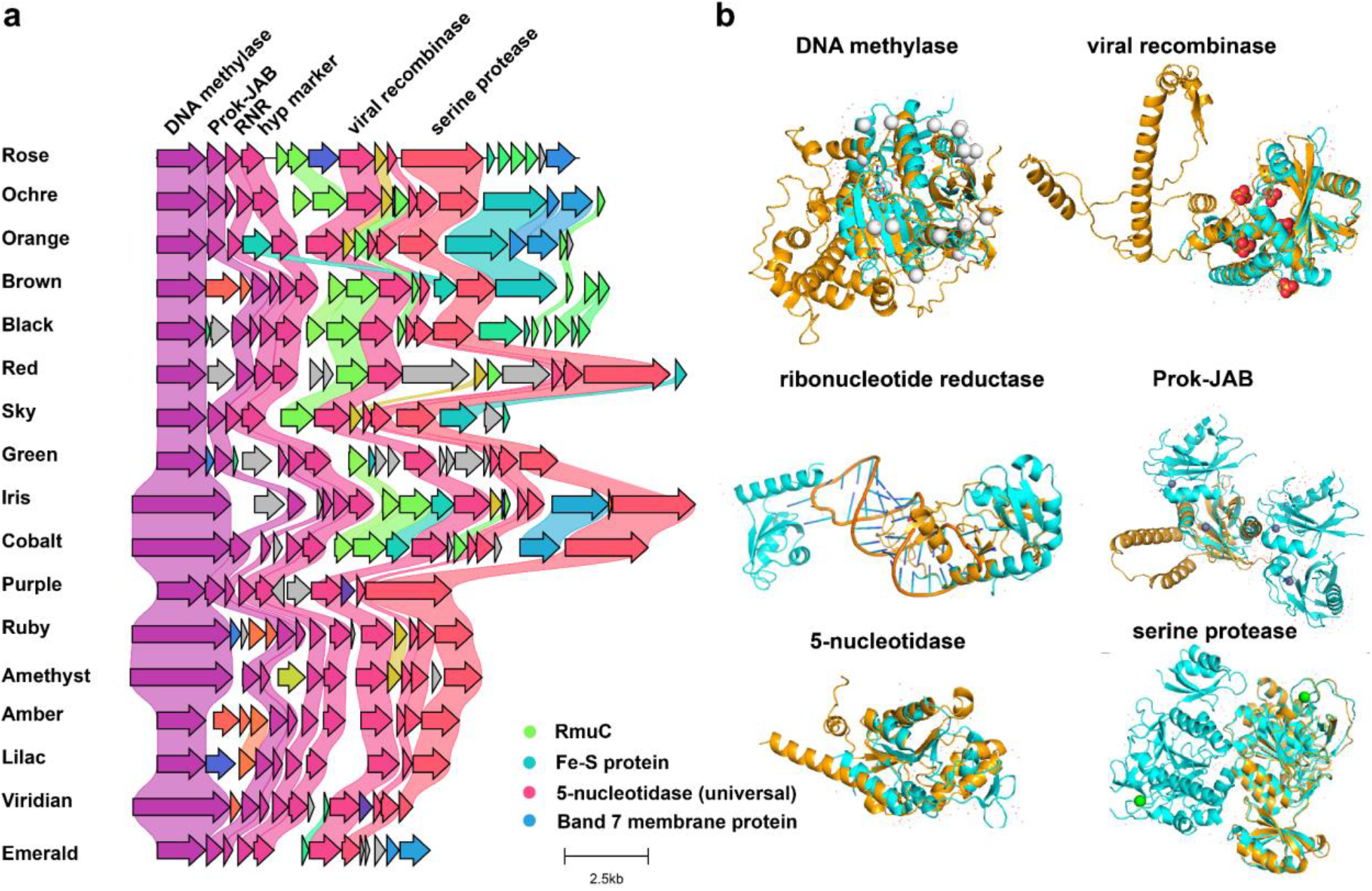
Conserved genomic region in all 17 Borgs comprises nucleotide processing machinery. **a**. Colored genes highlight homologs based on amino acid identity. The alignment was generated with clinker using the default settings. **b**. Protein structures from Orange Borg were predicted with AF2 and are shown in Orange. Best structural hits were obtained through PDBefold search. The best structural hit for DNA methylase is human NMRT-1 (PDB match 5e1b, RMSD = 2.81), for the viral recombinase is lambda exonuclease (PDB match 6m9k, RMSD = 1.84), for the ribonucleotide reductase (RNR) it is the RNA-binding XRRM domain of human LARP7 (PDB match 6d12, RMSD = 2.06), for the Prok-JAB it is AF2198, a JAB1/MPN domain protein from *Archaeoglobus fulgidus* (PDB match 1oi0, RMSD = 3.32), for the 5-nucleotidase it is a magnesium dependent phosphatase 1 (MDP-1; PDB match 1u7o, RMSD = 2.42), for the serine protease it is IS1-inserted Pro-subtilisin E (PDB match 3whi, RMSD = 1.03).

**Supplementary Figure 5.**
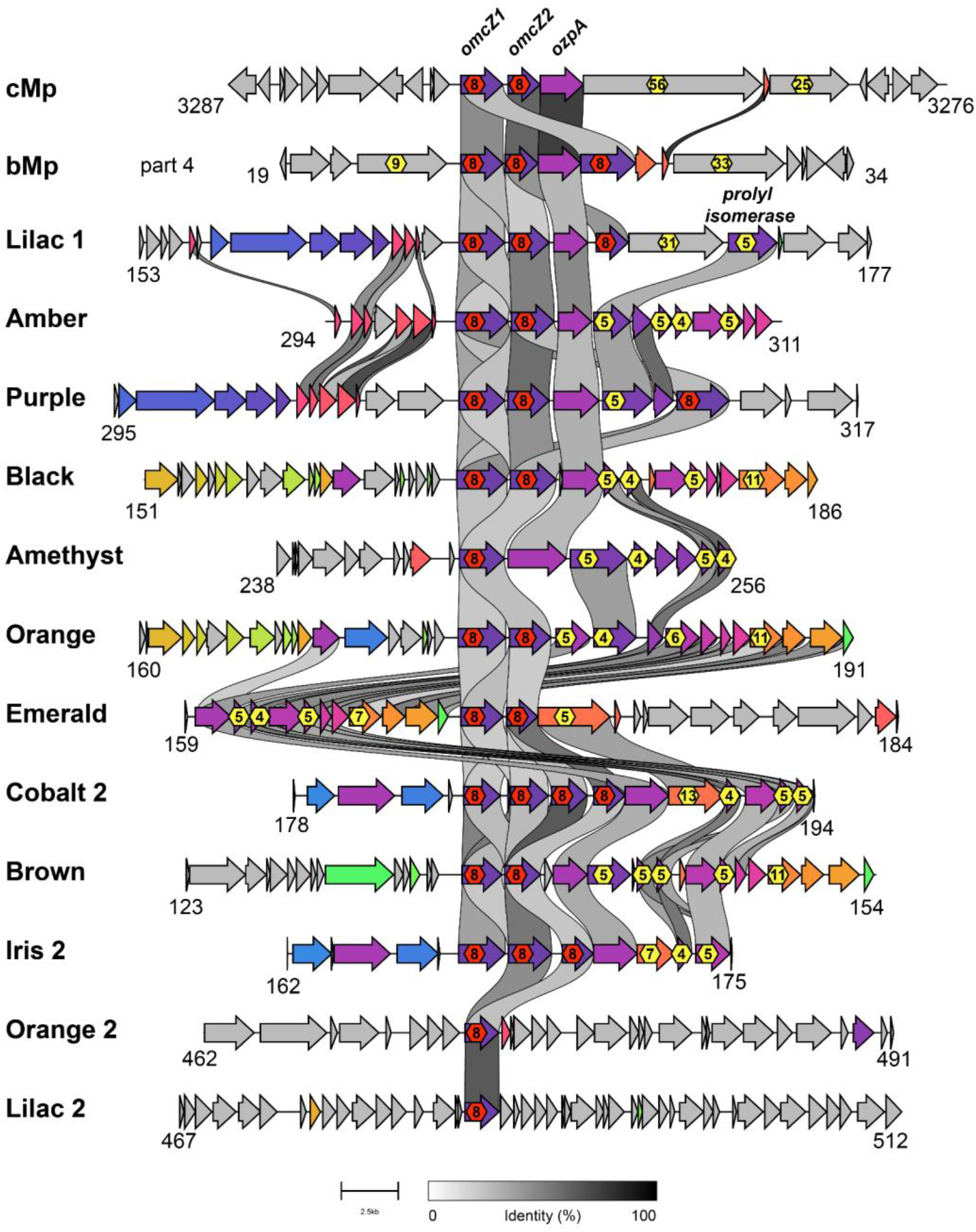
Genomic neighborhood alignments of potential nanowire-forming OmcZ clusters in Borgs and *Methanoperedens.* Alignments were performed with clinker and gene coordinates are shown.

## Supplementary Information [xlsx]

**Table S2. Annotations and features of all proteins encoded in the 17 Borgs.**

**Table S3. Protein subfamilies consistently encoded in Borgs.**

**Table S4. MHC proteins in Borgs, bMp and cMp. All proteins with >= 3 CxxCH motifs are listed here.**

**Table S5. Protein subfamilies established from the 17 Borgs, cMp and bMp.**

**Table S7. Normalized abundance of Borgs and *Methanoperedens* species across 70 samples.**

**Table S9. CheckV output of 17 Borg genomes.**

**Table S10. GeNomad output of 17 Borg genomes.**

**Table S11. Annotations and features of all proteins encoded in cMp and bMp.**

**Table S12. List of elements shown in the metabolic reconstruction of *Methanoperedens* and Borgs depicted in Figure 4**.

**Table S13a. Summary of metatranscriptomic analysis.**

**Table S13b. Number of features per genome.**

**Table S13c. Percentage of genes with transcript.**

**Table S14. Metatranscriptomic data of Borgs and *Methanoperedens* from four soil samples recovered from the wetland site in Lake County, CA.**

**Table S15. Coverage of Black Borg and bMp in nanopore metagenomic dataset originating from the same samples as the metatranscriptomic dataset.**

